# Fast and Deep Phosphoproteome Analysis with the Orbitrap Astral Mass Spectrometer

**DOI:** 10.1101/2023.11.21.568149

**Authors:** Noah M. Lancaster, Pavel Sinitcyn, Patrick Forny, Trenton M. Peters-Clarke, Caroline Fecher, Andrew J. Smith, Evgenia Shishkova, Tabiwang N. Arrey, Anna Pashkova, Margaret Lea Robinson, Nicholas Arp, Jing Fan, Juli Hansen, Andrea Galmozzi, Lia R. Serrano, Julie Rojas, Audrey P Gasch, Michael S. Westphall, Hamish Stewart, Christian Hock, Eugen Damoc, David J. Pagliarini, Vlad Zabrouskov, Joshua J. Coon

## Abstract

Owing to its roles in cellular signal transduction, protein phosphorylation plays critical roles in myriad cell processes. That said, detecting and quantifying protein phosphorylation has remained a challenge. We describe the use of a novel mass spectrometer (Orbitrap Astral) coupled with data-independent acquisition (DIA) to achieve rapid and deep analysis of human and mouse phosphoproteomes. With this method we map approximately 30,000 unique human phosphorylation sites within a half-hour of data collection. The technology was benchmarked to other state-of-the-art MS platforms using both synthetic peptide standards and with EGF-stimulated HeLa cells. We applied this approach to generate a phosphoproteome multi-tissue atlas of the mouse. Altogether, we detected 81,120 unique phosphorylation sites within 12 hours of measurement. With this unique dataset, we examine the sequence, structural, and kinase specificity context of protein phosphorylation. Finally, we highlight the discovery potential of this resource with multiple examples of novel phosphorylation events relevant to mitochondrial and brain biology.

## INTRODUCTION

Protein phosphorylation is an essential post-translational regulatory mechanism for myriad cellular functions including apoptosis, inflammation, metabolism, proliferation, protein trafficking, and many others^1^. Global detection of which proteins, and perhaps most importantly which residues, are subject to this dynamic modification has been a key technological gap for decades.^2^ In the early 2000s, key advancements in technologies to enrich phosphorylated peptides – prior to mass spectrometry (MS) analysis – enabled a new era of large-scale phosphorylation discovery experiments.^3–13^ The evolution of that work, combined with gradual, but steady, improvements in both MS hardware and data analysis tools make it possible to map and quantify thousands of phosphorylation sites in a single experiment.^14–27^ Still, current phosphoproteomic analyses are neither routine nor straightforward when compared to protein detection and quantification.

Phosphoproteome analysis remains challenging due to four main requirements: need for site localization, dynamic range, reproducibility, and throughput. The quality of the tandem mass spectra required to determine precisely upon which residue the phosphoryl group resides is higher than that required to identify an unmodified peptide. Alternative dissociation methods, such as electron transfer dissociation (ETD), and improved mass resolving power and accuracy are both strategies that can aid in improving spectral quality and increasing the likelihood that the detected site can be localized with high confidence.^15,21,28,29^ Additionally, sensitivity and dynamic range of the mass analyzer can also be important for detection of low level, but critical, product ions.^11,23,27^ Next, the often sub-stoichiometric amounts of phosphorylated protein itself elevate the difficultly.^2,13,17–19^ Beyond that, previous studies demonstrate that this dynamic range problem is further exacerbated in that ∼10% of detected phosphopeptides account for ∼80% of the observed signal.^30^ This tremendous dynamic range necessitates both enrichment of phosphorylated peptides and chromatographic fractionation ahead of conventional capillary liquid chromatography tandem MS (nLC-MS/MS).^2,17,18^

Phosphorylation site quantification is often essential to elucidating biological insight.^17^ The requirements outlined above also add challenges to achieving this goal. Quantification of a protein, for example, is done by summing the signals of multiple unique peptides all stemming from that single protein. Thus, if a single peptide is not reproducibly detected, the overall measurement can still be made. But unique phosphopeptides cannot be summed with other signals; they must be reproducibly and reliably detected from one sample to the next. Such demands make performing truly large-scale (i.e., > 100 samples) comparisons of phosphoproteomes very difficult.^18^ Finally, getting sufficient depth to detect targets of a particular kinase, for example, might require extensive fractionation, as discussed. However, scaling that experiment to multiple conditions or samples is often not possible from a throughput perspective, despite all the caveats noted. We conclude that global and quantitative phosphoproteomics technologies require improvement to enable routine and truly large-scale phosphoproteome measurements.

Recently, a new type of mass analyzer has been described – the Asymmetric Track Lossless analyzer (Astral^TM^). The Astral analyzer can achieve high resolving powers (∼80,000) and mass accuracy (5 ppm), single ion detection limit, and MS/MS scan speeds up to 200 Hz.^31,32^ Here we describe the use of a quadrupole-Orbitrap^TM^-Astral hybrid MS instrument for the analysis of phosphopeptides. Specifically, we examine the ability of this system to perform MS/MS scans of complex mixtures of phosphopeptides separated over durations ranging from 7 to 60 minutes. We examine the performance of these methods for various peptide mass loads and data acquisition settings as a function of detected and localized phosphorylation sites and overall reproducibility. Validation of site localization and quantification was accomplished using synthetic phosphopeptide standards spiked into a yeast phosphopeptide background. Next, we benchmarked the performance of our Orbitrap Astral method against an Orbitrap Ascend and a previously described timsTOF Pro method by replicating an EGF stimulation of HeLa cells.^33^ Finally, we leverage this technology to collect an atlas of phosphorylation in the mouse – generating tens of thousands of localized phosphorylation sites from each of twelve unique tissues. With these data we present, to our knowledge, the deepest mouse phosphoproteome collected in a single study. Using this atlas combined with AlphaFold predicted protein structure^34^, we confirm existing hypotheses that that most phosphorylation events are directed towards unstructured regions of proteins.^35–37^ The incorporation of a previous kinome atlas allows investigation of tissue-specific kinase activity.^38^ We additionally provide examples of how our novel resource can be mined for key phosphorylation events on biologically relevant proteins

## RESULTS

### Rapid phosphopeptide analysis with the Orbitrap Astral mass spectrometer

Owing to its extremely fast MS/MS scan rate and high sensitivity, we hypothesized that the Orbitrap Astral^TM^ mass spectrometer could resolve many of the aforementioned challenges in analyzing phosphoproteomes. Specifically, the Orbitrap Astral MS comprises a conventional quadrupole-Orbitrap coupled with a new mass analyzer (Astral, **Figure 1A**). The combination of low ion losses and single ion detection drives the high sensitivity of the Astral analyzer, while very fast MS/MS scan rates allow it to cycle through large numbers of targets. In a typical Orbitrap Astral method, the Astral analyzer is set to generate 200 MS/MS spectra per second while the Orbitrap analyzer in parallel generates slower high resolution and high dynamic range MS data.^31^

**Figure 1.**
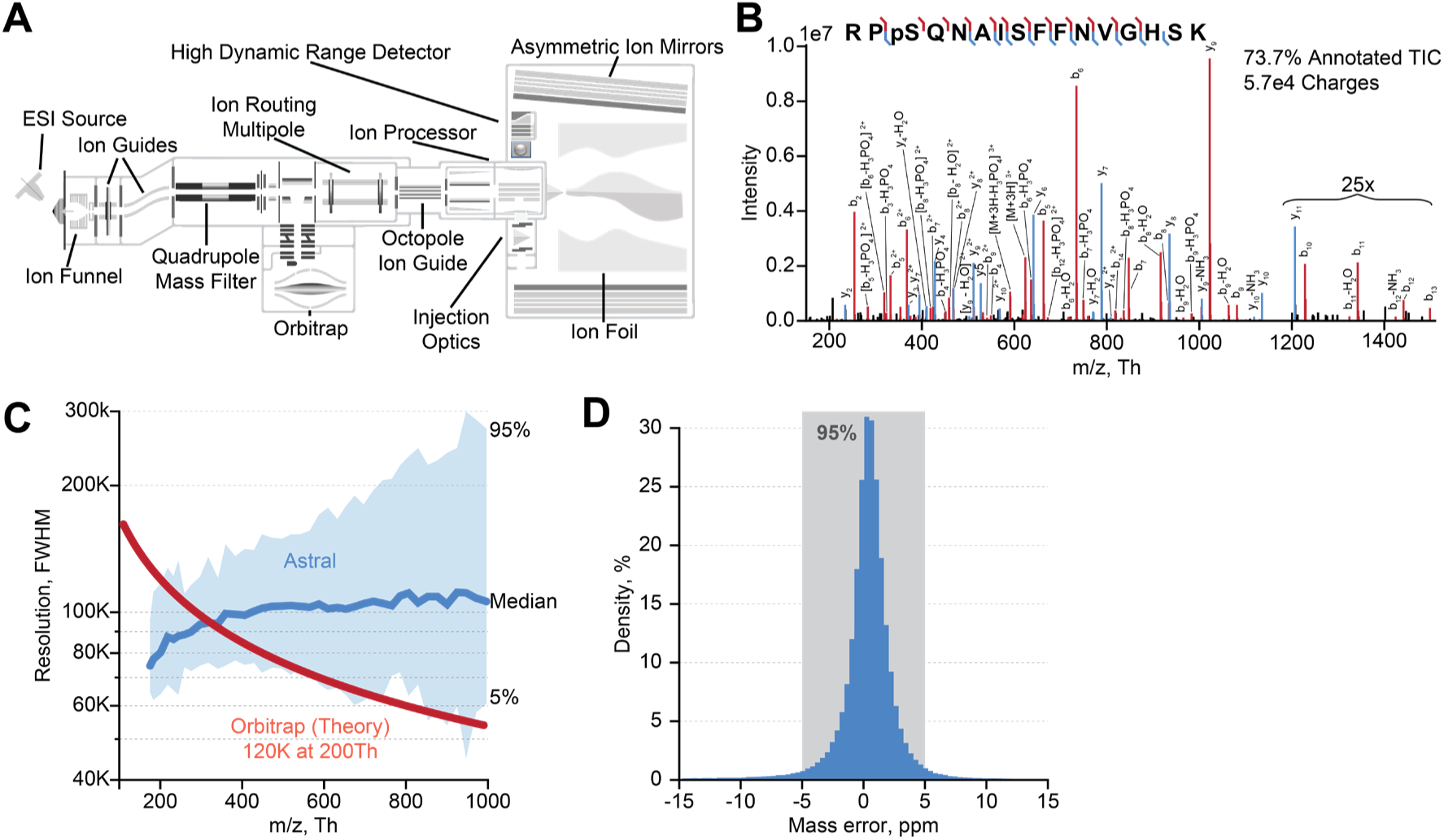
Overview of Orbitrap Astral MS and its key figures of merit. (A) Orbitrap Astral instrument schematic, highlighting the quadrupole, Orbitrap, and Astral analyzers. (B) Tandem mass spectrum of representative phosphopeptide collected using the Astral analyzer. (C) Distribution of Astral analyzer resolving power as a function of mass from phosphopeptide product ion spectra collected here. (D) Phosphopeptide product ion mass measurement error from Astral analyzer.

To test the utility of the Orbitrap Astral MS for the analysis of phosphopeptides, we purified tryptic phosphopeptides from HEK239T cells, loaded them onto a nanoflow capillary column with a pulled electrospray emitter^39^, and eluted them into the Orbitrap Astral mass spectrometer. For this initial experiment, we utilized a data-dependent acquisition (DDA) method wherein MS scans were acquired in the Orbitrap analyzer while MS/MS scans were collected using the Astral analyzer. **Figure 1B** represents an example single-scan tandem mass spectrum that was identified following a traditional MaxQuant database search.^25,40^ Here, a triply-protonated precursor having *m/z* value of 623.6314 and a sequence of RPsQNAISFFNVGHSK was selected, dissociated with beam-type collisional activation (HCD), and analyzed in the Astral analyzer– all within 5 ms. In total, from this single 30-minute nLC-MS/MS DDA experiment, we collected 3,201 MS and 174,944 MS/MS scans, from which we identified 12,327 phosphopeptides corresponding to 9,537 unique sites of phosphorylation. Note a previous study using the Orbitrap Tribrid platform with a DDA method reported a depth of ∼9,500 phosphosites following a 120-minute method.^20^ These MS/MS scans were collected within ∼6.3 ms on average, enabling extremely fast sampling of selected precursors, and sometimes exhausting the precursors available to sequence. Next, we investigated the Astral analyzer’s capabilities by plotting the measured resolving power (**Figure 1C**) and mass accuracy (**Figure 1D**) for these product ions. The upper bound of observed resolving powers likely correspond to peaks with low numbers of ions, whereas the lower bound presumably arises from high intensity peaks exhibiting space charging.^31^ Note the Astral resolving power persists with increasing *m/z* value. Finally, mass accuracy, as measured for these product ions, was within 5 ppm for 95% of the measurements.

With the exceptional speed of the Astral mass analyzer, combined with its ability to deliver high mass resolution and accuracy, we next wondered how this instrument would perform for DIA analysis of phosphopeptides.^33,41–45^ Accordingly, we analyzed the same tryptic phosphopeptide mixture using a DIA method. The speed of the Astral analyzer allowed for use of DIA windows as narrow as 2 *m/z* across the range of 380 to 980 *m/z* while still maintaining a cycle time of 1.5 seconds. This scan method is visualized in **Figure 2A** with the high-resolution Orbitrap MS scans (240K resolving power) denoted in blue (∼0.6 s cycle time) and the Astral MS/MS scans depicted in red.

**Figure 2.**
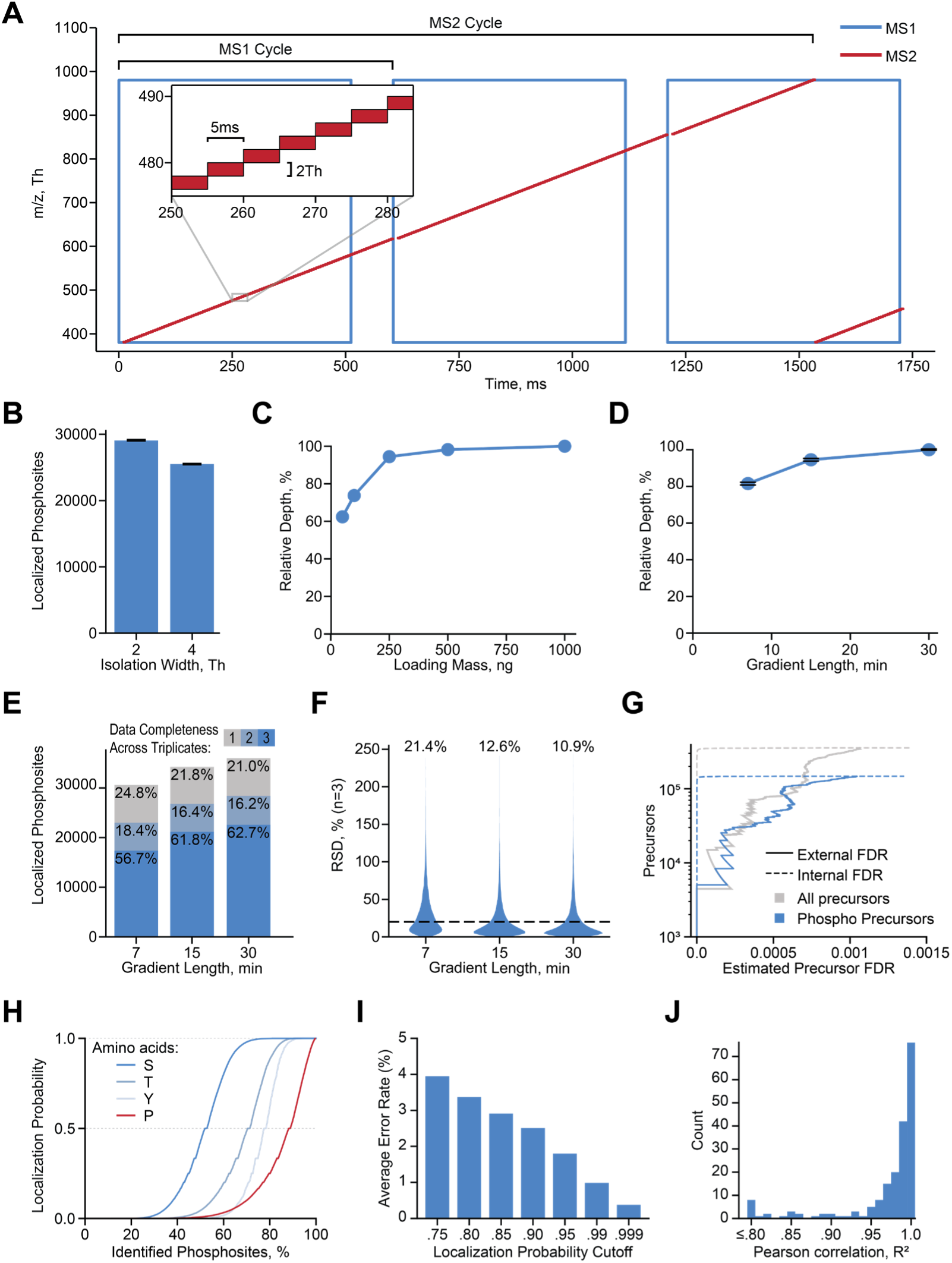
Overview of DIA phosphoproteomics on the Orbitrap Astral MS. (A) Illustration of DIA acquisition scheme. (B) Number of localized phosphorylation sites identified using DIA method with various isolation widths. (C) Evaluation of phosphopeptide loading mass on performance. Note data was collected using a 30-minute active gradient and 2 *m/z* DIA isolation width. (D) Evaluation of gradient length on performance. (E) Evaluation of phosphosite identification reproducibility (note this plot uses results from (D)). (F) Evaluation of phosphosite quantitative precision. Here the relative standard deviation of phosphosite quantities is shown for phosphosites detected across triplicate injections with the median value displayed for each active gradient length. (G) Comparison of External (maize/human entrapment experiment) and Internal (Spectronaut) FDR on precursor level. (H) Phosphoproline Decoy Search to test reliability of localization algorithm. The cumulative distribution of localization probabilities is shown for phosphorylation events at different amino acids. (I) The average site localization error rate as a function of localization probability cutoff is shown. Synthetic phosphopeptide standards spiked into a yeast phosphopeptide background were used as a ground truth for error rate determination. (J) The distribution of R^2^ values for linear calibrations curves are shown for phosphopeptides standards detected in at least three concentrations across a five-point dilution series into a constant yeast phosphopeptide background.

A 250-ng injection of tryptic phosphopeptides was analyzed with a 30-minute nLC separation and the DIA method described above (see **Supplemental Figure 1A** for the base peak chromatogram). The resultant data was searched both with Spectronaut and a developmental build of CHIMERYS (see Methods). From the Spectronaut search, we localized 29,190 phosphorylation sites (**Figure 2B**, see **Supplementary Figure 2A** for comparison of Spectronaut and CHIMERYS, and **Supplementary Figure 2B and 2C** phosphopeptide multiplicity and localization probability distributions, respectively). We note that the use of a wider DIA window (4 *m/z*) produced 14% fewer localized phosphorylation sites. Next, we examined the impact of loading amount on performance using a dilution series ranging from 1 µg to 50 ng loaded on column (**Figure 2C**). We note that similar results are obtained down to 250 ng loads. The sensitivity of the system is evident in that a 50-ng load produced ∼60% of the detected sites observed with a 1 µg load. To determine whether the Astral scan speed would allow for increased throughput, we next evaluated a range of active gradient lengths (**Figure 2D, Supplementary Figure 1A**). A four-fold reduction in gradient length (30 to 7 minutes) results in just a ∼20% reduction in localized phosphorylation sites. Additional method parameters are justified in the **Supplemental Text**.

To determine the impact of gradient length on reproducibility, we performed two additional analyses. First, we examined identification reproducibility across triplicate injections (**Figure 2E**). The 7-, 15-, and 30-minute active gradient methods resulted in 56.7%, 61.8%, and 62.7% of phosphosites detected across all three injection replicates, respectively. Second, we assessed quantitative precision by determining percent relative standard deviation (RSD) of phosphorylation sites detected across all three replicates (**Figure 2F**). The 15– and 30-minute active gradients achieved median RSDs of 10.9 and 12.6%. Not unexpectedly, the 7-minute active gradient method has considerably reduced precision (21.4% median RSD), most likely due to insufficient sampling across the narrower elution peaks (**Supplementary Figure 1B-D**).

Since these data were collected with a novel mass analyzer and new instrument, we next sought to ensure that the phosphosites were identified and localized accurately. To test this, we first performed an entrapment search with a combined human and maize protein database.^46^ In this experiment, which is commonly used to evaluate internal false discovery rate (FDR), maize false identifications allow for the calculation of an external FDR. The results of this analysis are shown in **Figure 2G**, where internal and external FDR is plotted for all precursors and phosphopeptide precursors. For lower q-values the internal and external FDRs disagree; however, these curves converge at approximately the 1% FDR threshold, suggesting that the identifications reported here are reliable. We conclude that, as the DIA software tools continue to evolve and are adopted for the new data type, this discordance will be diminished.

In the case of phosphoproteomics, peptide identification is not the final result – the site of phosphorylation is ideally localized to a specific residue with confidence. To evaluate the quality of phosphosite localization, we searched these data with phosphorylated proline as a variable modification where any detected proline phosphorylation is false.^47^ S, T, and Y all have a significant amount of highly confident localized phosphosites whereas the P does not (**Figure 2H, Supplemental Figure 2H**). To further explore localization confidence, we analyzed a set of synthetic phosphopeptide standards that had been spiked into a complex mixture yeast tryptic phosphopeptides at various concentrations. Note this same 225 synthetic phosphopeptide mix has been used by others and thereby provides a means of both assessment and comparison^41,48^; specifically, the experiment allows for calculation of localization error rate based on ground truth knowledge. From these data we conclude that, depending on localization probability cutoff, the error rate ranges between 1 and 5 percent, a similar trend as the previous study (**Figure 2I, Supplemental Figure 3A**).^41^ **Supplemental Figure 3B** demonstrates that we retain the majority of correct precursors even at more stringent localization probability cutoffs. Taken together, these results confirm that a library-free DIA search, using MS/MS spectra from an Astral analyzer, can reliably detect and localize phosphorylation sites.

The aforementioned phosphopeptide standards also provided an opportunity to evaluate quantitative linearity.^42,48^ To accomplish this, phosphopeptide standards were diluted from 10,000 to 39 attomole (per standard, on-column) into a constant complex mixture of yeast tryptic phosphopeptides. Using the 15-minute DIA Orbitrap Astral method, we analyzed the various spike-in samples and then calculated the linear fit between the observed MS intensities and the load amount. **Supplemental Figure 3D** displays three randomly chosen curves while **Figure 2J** confirms that the majority of the phosphosite calibration curves have R^2^ values greater than 0.95, indicating good quantitative linearity. **Supplemental Figure 3C** confirms that even at loads as low as 39 attomole, half of the phosphorylation sites were detected across all three injections. Finally, we note that the phosphopeptide standard intensity distributions observed across the dilution series overlap with the yeast phosphopeptide intensity distribution (**Supplemental Figure 3E**). Together, these results demonstrate that our method exhibits reliable detection and quantification over the concentration range typically observed in phosphoproteomic analysis.

### Comparison of Orbitrap Astral to other phosphoproteomics platforms

To compare the performance of the Orbitrap Astral to other phosphoproteomics platforms we replicated the HeLa EGF stimulation experiment as previously reported by Skowronek et al. using the timsTOF Pro hybrid mass spectrometer with dia-PASEF acquisition.^33^ Briefly, we cultured HeLa cells in triplicate followed by 15 minutes of EGF stimulation, cell lysis, protein extraction, trypsin digestion, and phosphopeptide enrichment. The resulting phosphopeptide samples were then analyzed with a 15-minute active gradient, closely replicating the total acquisition time for the Skowronek et al. publication, with detection using either the Orbitrap Astral or Orbitrap Ascend^TM^ Tribrid mass spectrometers. To enable a direct comparison, we searched our raw files and those from the Skowronek et al. study with the same protein database and Spectronaut library-free method. **Figure 3A** shows the phosphoproteomic depth and completeness across triplicates for all three datasets. The Orbitrap Astral method provided the deepest phosphoproteomic analysis and yielded near 60% phosphosite completeness across replicates from approximately three-fold more localized sites as compared to either the timsTOF Pro or Orbitrap Ascend instruments. Note the Orbitrap Ascend is not optimally operated in DIA mode given its slower scan speed; however, we chose to use a DIA method for direct comparison. Interestingly, the Orbitrap Ascend generated the highest level of phosphosite overlap of all three instruments at 64%.

**Figure 3.**
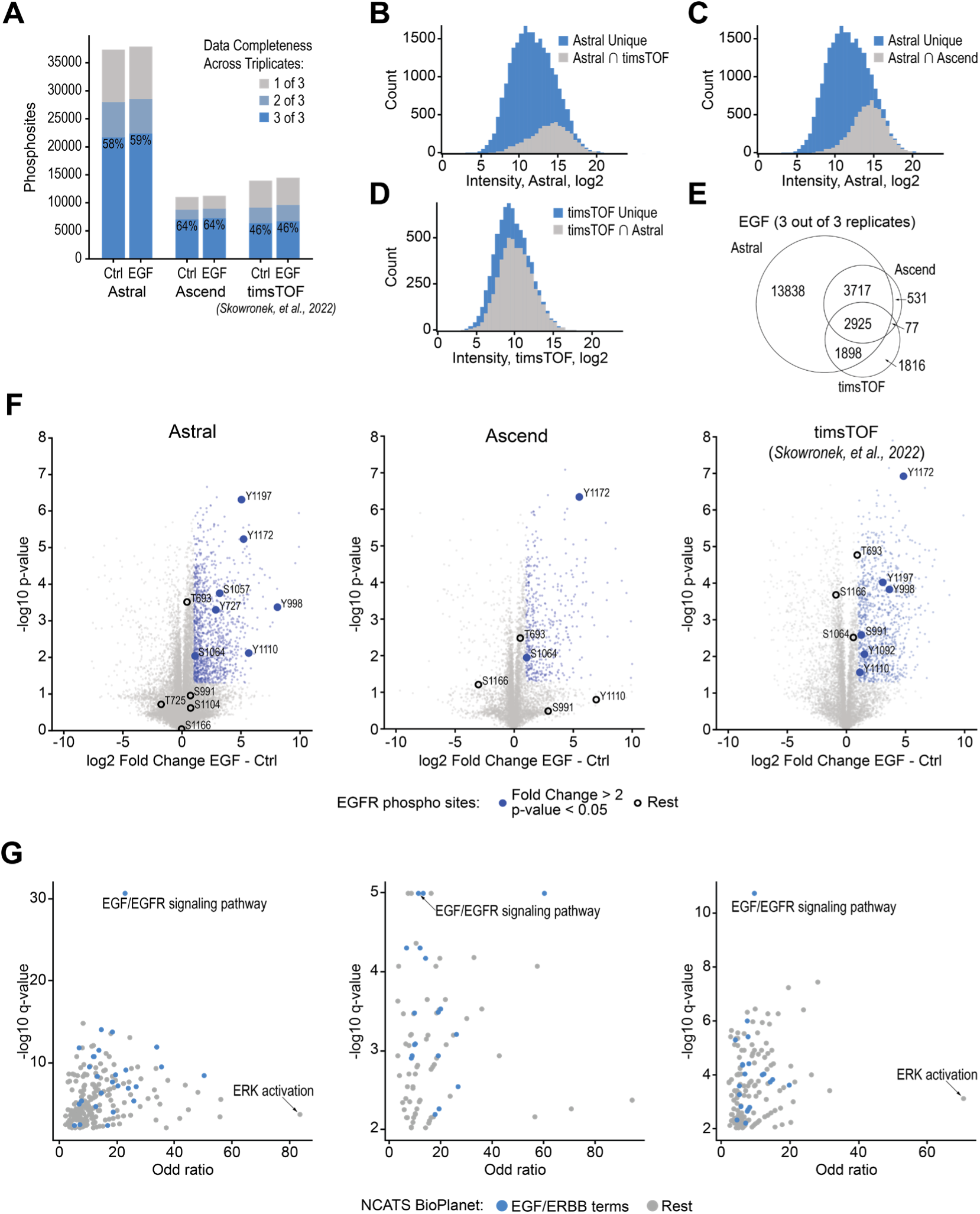
Biological validation of phosphoproteomics platforms using EGF stimulation. (A) Reproducibility of detected phosphosites across biological triplicates for three phosphoproteomics platforms – Orbitrap Astral, Orbitrap Ascend, and timsTOF Pro.^33^ Intensity distribution of phosphosites consistently detected (3 out of 3 EGF stimulated replicates) by Orbitrap Astral with overlap indicated for (B) timsTOF Pro and (C) Orbitrap Ascend. (D) Intensity distribution of phosphosites consistently detected (3 out of 3 EGF stimulated replicates) by timsTOF Pro with overlap indicated for Orbitrap Astral. (E) Venn diagram of phosphosites detected across all three mass spectrometry platforms. (F) Vulcano plots of phosphosites between EGF stimulation and control across platforms with phosphosites meeting the differential expression criteria (fold-change >2, p-value < 0.05) indicated in blue and EGFR phosphosites labeled. (G) Pathway enrichment analysis using the NCATS BioPlanet terms was performed for the three platforms with EGF/ERBB-associated terms indicated in blue.

To examine the dynamic range of detected phosphorylation sites across platforms we plotted the intensity distributions of phosphorylation sites detected in all three replicates using only the EGF-stimulated data. **Figure 3B** and **3C** present the Orbitrap Astral intensity distributions with overlapping sites indicated for either the timsTOF Pro or Orbitrap Ascend datasets, respectively. Both plots reveal that the unique phosphosites in the Orbitrap Astral dataset are biased toward lower intensities. In contrast, unique phosphosites for the timsTOF Pro dataset are distributed more evenly across the intensity range (**Figure 3D**). These observations suggest that our Orbitrap Astral exhibits improved sensitivity, resulting in a larger dynamic range. Note the Orbitrap Astral and Ascend shows higher overlap as compared to the timsTOF Pro (**Figure 3E**); we suppose this difference is likely due to variation in EGF treatment and sample preparation.

To see how this performance variation translates to biological discovery we performed a differential expression analysis between control and EGF-stimulated samples for all three datasets (**Figure 3F**). Each datapoint in these plots presents a phosphorylation site with the differentially expressed ones highlighted in blue. Phosphorylation sites occurring directly on the EGFR are labeled, many of which are detected across all platforms. To further compare we performed a pathway enrichment analysis from each dataset (**Figure 3G**) – all three platforms had the EGF/EGFR signaling pathway as the top enriched term. We conclude that despite differences in treatment, sample preparation, and instrumentation, all technologies could arrive at the same biological conclusions.

### Deep phosphoproteome analysis of the mouse

Having demonstrated the speed and sensitivity of the Orbitrap Astral MS for reliable identification of phosphopeptides, we next sought to leverage this methodology to generate a comprehensive phosphoprotein atlas of the mouse. In 2010, Huttlin and co-workers described the first mouse phosphoproteome atlas, reporting 35,965 phosphorylation sites detected following 10 days of mass spectrometry analysis and serving as a valuable reference point for this study.^49^ **Figure 4A** presents our overall experimental design wherein proteins were isolated, digested, and enriched for phosphorylation from 12 distinct tissues. For each tissue, the purified phosphopeptides were separated offline and concatenated into four fractions, each of which were analyzed using a 15-minute DIA nLC-MS/MS method on the Orbitrap Astral MS. In total, this experiment required 12 hours of active data acquisition time and resulted in the detection of 81,120 unique phosphorylation sites (**Supplementary Table 3**). Note to provide complementary protein abundance we separately analyzed non-enriched peptides from each tissue (**Supplementary Figure 4**, **Supplementary Table 4**).

**Figure 4.**
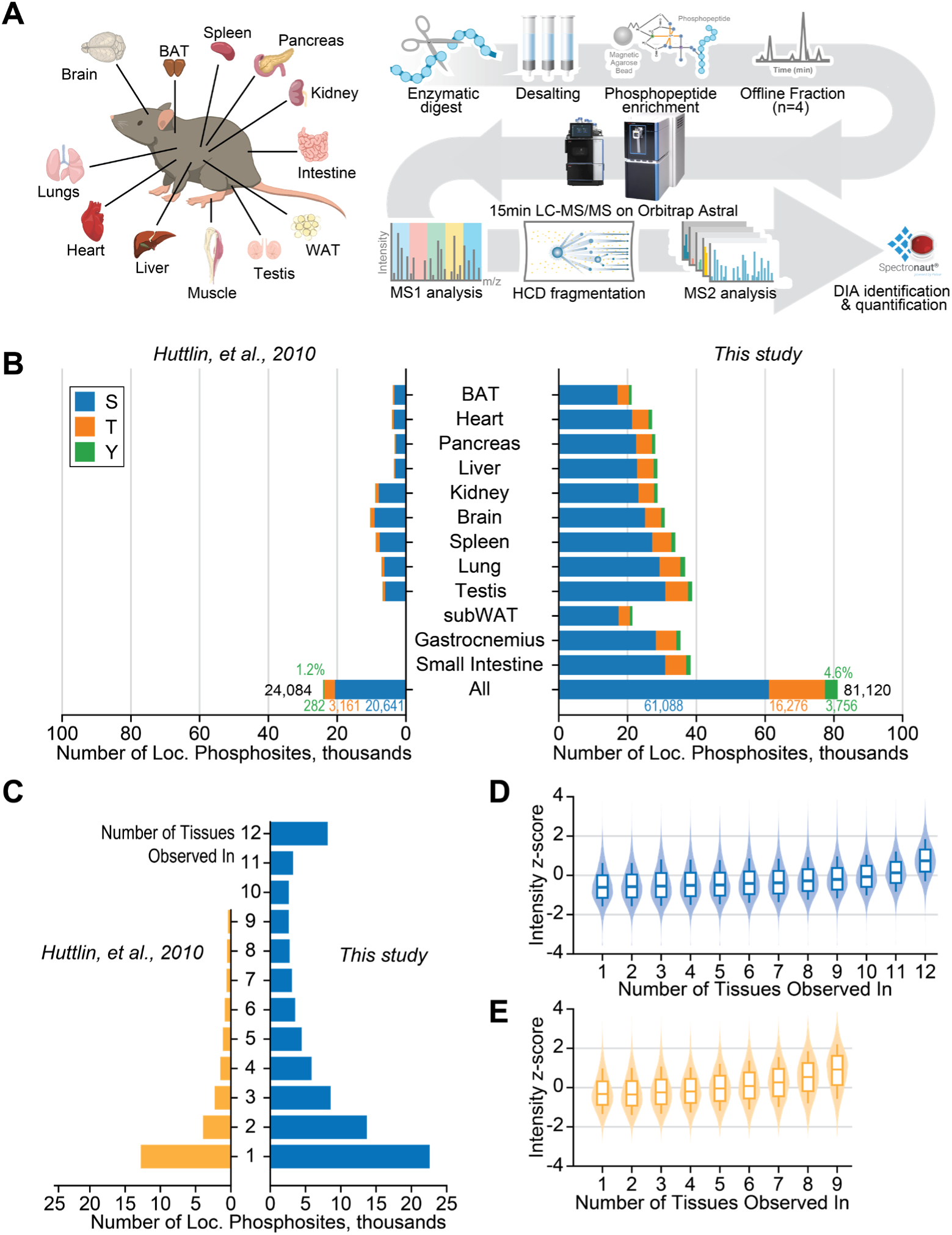
Mouse Phosphorylation Atlas Workflow and Results. (A) Mouse Phosphorylation Atlas Workflow. (B) Result for Mouse Tissue Phosphoproteomic Analysis. Numbers of unique phosphorylation sites are shown for each tissue and the total unique sites with the fraction of S,T, and Y localizations indicated. The Huttlin et al. results were generated by researching the raw data in MaxQuant using the same protein database used in this study. (C) Tissue Specificity of Detected Phosphorylation sites. The y-axis indicates the number of tissues in which a phosphosite was detected. (D) and (E) Intensity distributions for phosphorylation sites detected in a given number of tissues for this study and by Huttlin et al., respectively.

Depending upon the source tissue, the number of identified phosphorylation sites ranged from approximately 20,000 to 40,000 sites (**Figure 4B**). Of the 81,120 unique sites identified, 61,088 were localized to S; 16,276 localized to T; and 3,756 to Y. It is noteworthy that ∼4.6% of the total sites identified stem from pY. Previous studies have placed this number closer to ∼2% and often rely on pY-specific antibodies for enrichment of this perhaps most functionally important phosphorylation site.^30,50^ We suppose that the increased sensitivity and depth afforded by this new analyzer permits the detection of these low-expression phosphorylation events (**Supplementary Figure 5C**). Also shown in **Figure 4B** are the results, by tissue, from the previous mouse atlas. Note these raw data were reprocessed using MaxQuant with the matching protein database and filtered using similar quality metrics. On a per tissue comparison, the method leveraged here provides a 5-fold boost in the number of localized phosphorylation sites in approximately 1/24^th^ the time.

Next, we plotted the distribution, by tissue, of phosphorylation sites and whether they were detected in both studies (**Supplementary Figure 5A**). Despite the age and strain differences between mice, and the dynamic nature of phosphorylation, we see good overlap in detection of sites (median 55.3%). Tissue-specificity was also plotted (**Figure 4C** and **Supplementary Figure 5B**) and compared to Huttlin et al. In both studies, a large portion of phosphorylation sites appear to have single tissue specificity; however, our results indicate that a sizeable number of sites are detected across all twelve tissues. This result contrasts with the Huttlin work and likely stems from the increased sensitivity and reproducibility afforded by the new analyzer and DIA method. With this result, we expect that many sites may indeed be present across all tissues and that further improvements in sensitivity and reproducibility are likely needed to detect them. This hypothesis is further evinced by the results shown in **Figure 4D-E**, where we examined the detected peak intensities for each site as a function of how many tissues in which it was observed. From these data, both studies show concordance between number of tissues detected and overall signal strength (*i.e.*, abundance).

Furthermore, we compared our comprehensive phosphoprotein atlas of the mouse with the most recent multi-tissue study conducted by Giansanti and co-workers.^51^ Similar to our sample preparation, the phosphopeptides from multiple tissues were offline fractionated and further concatenated into 4 samples. These samples were measured using a 90-minute gradient DDA on the Q Exactive Orbitrap HF MS system. Despite a 6-fold shorter measurement time, the Orbitrap Astral detected 2-3 times more phosphosites per tissue, with improved agreement with our dataset compared to Huttlin et al. (median 64.2% versus 55.3%; **Supplementary Figure 6A**). Interestingly, Giansanti’s study shows a similar trend towards a higher number of sites that are present across all tissues (**Supplementary Figure 6B**). The portion of pY in this study is 1.1% (**Supplementary Figure 6C**), which additionally highlights the capability of combining Astral sensitivity with DIA acquisition to enable the detection of 4.6% pY in our data.

### Phosphorylation sites in the context of protein sequence and structure

Phosphorylation sites within proteins exhibit a specificity determined by both the sequence and structural motifs. In our investigation, we sought to unravel this specificity by clustering phosphorylation sites and their flanking regions. To achieve this, we computed all pairwise comparisons of phosphosite sequences (site and five amino acids in both C and N-terminal direction) and projected this comparison onto a 2D plane using the t-SNE algorithm (**Supplementary Figure 7A**). Notably, our analysis highlighted the importance of the phosphorylation amino acid as a major factor driving separation (**Supplementary Figure 7B**). That is clusters 1-8 are Ser-based, while 9 is Tyr-based, and 10-12 are Thr-based. Intriguingly, upon repeating the analysis while excluding the phosphorylated amino acid from binary comparisons (**Figure 5A**), we identified predominant clusters organized by specific motifs (**Figure 5B**), such as S/T-P (cluster 1), and RXX-S/T (cluster 9). Proline-directed protein kinases demonstrate a preference for phosphorylating serine or threonine residues immediately preceding a proline residue in proteins. Prominent examples of such kinases include extracellular signal-regulated kinases (ERKs) and cyclin-dependent kinases (CDKs). Leveraging annotations from the PhosphoSitePlus database^52^, our analysis revealed that the majority of ERK/CDK annotated phosphorylation sites belong to the S/T-P cluster (**Supplementary Figure 7C**). Note there is limited representation of Y in clusters 1 and 9, even though the phosphorylated residue was not considered in this analysis (**Supplementary Figure 7D**). These data confirm known biology that Y is not a substrate for P and R-directed protein kinases.^53^

**Figure 5.**
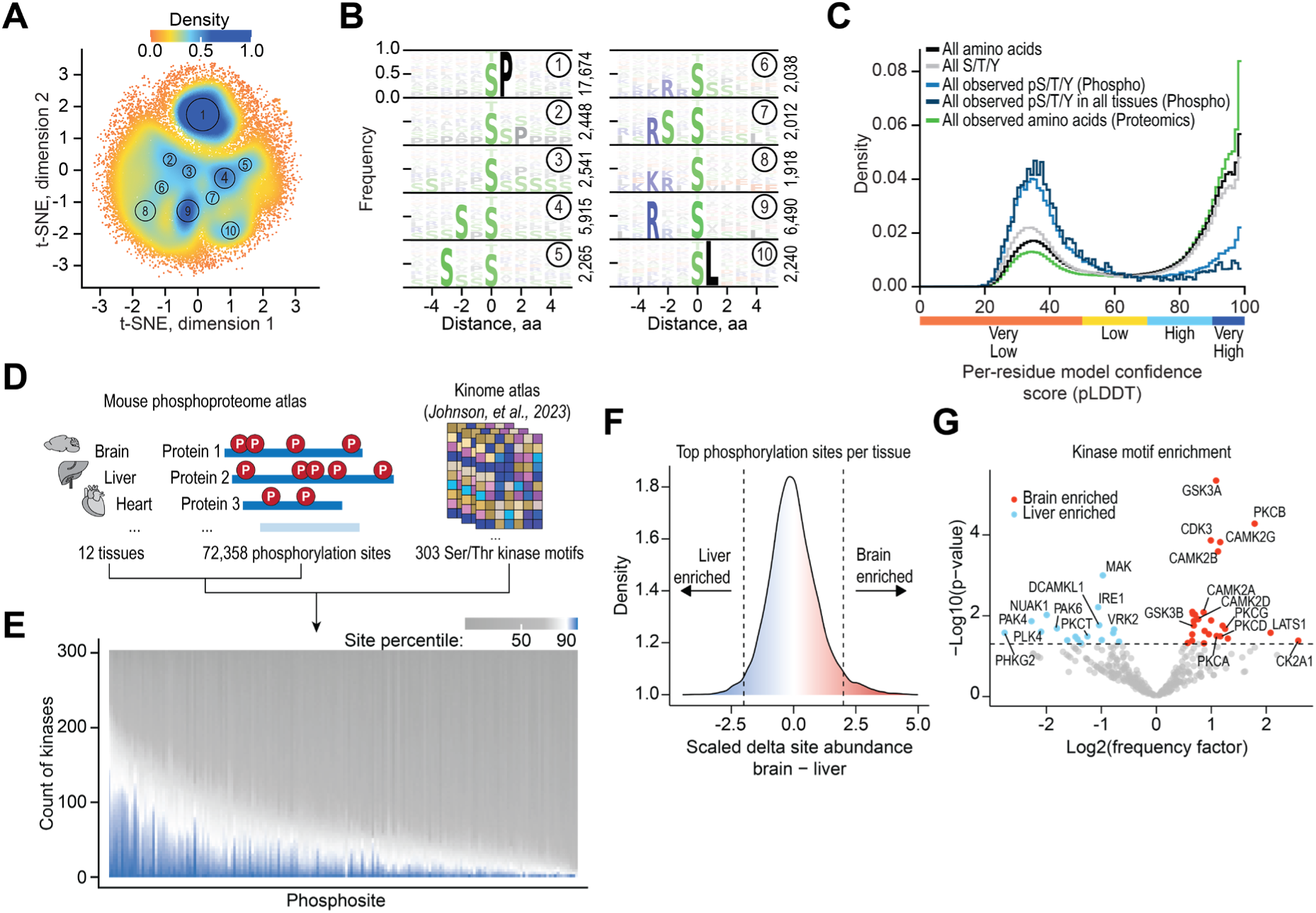
Sequence, Structural and Kinase Specificity Context of Phosphosites. (A) Two-dimensional representation of all phosphosites and their 5-amino acid flanking sequences, excluding the central amino acid from the comparison. Each cluster has been manually selected to emphasize the densest regions. (B) Sequence logo plot for all clusters depicted in (A). (C) Distribution of confidence scores for all amino acids, specifically S/T/Y, and for phosphorylated S/T/Y detected across all tissues. (D) Our mouse phosphoproteome data derived from nine tissues was applied to the kinome atlas search tool. (E), All phosphorylation sites detected in our study are plotted on the x-axis, sorted by the number of kinases that scored higher than 90 for a specific site. (F), Z-score transformed difference between abundances of shared phosphorylation sites in brain and liver tissue. Vertical dashed lines indicate thresholds for selection of phosphorylation sites that are used for kinase motif enrichment analysis. (G), Based on the top sites per tissue, a motif enrichment analysis was performed and the resulting frequency of how often a kinase was predicted to act on a site was plotted on the x-axis, along with the p-value on the y-axis. The scheme and types of analyses have been adapted from Reference 38.

To explore structural motifs, we mapped the 81,120 phosphorylation sites detected here onto protein structures predicted by AlphaFold.^34^ The UniProt structural library (assembled using AlphaFold) contains structures with varying degrees of certainty as measured by a per residue confidence score (pLDDT) where 0 is very low and 100 is very high confidence. Note that lower confidence scores are indicative of either flexible or disordered protein regions.^54^ Upon plotting the confidence score distribution for phosphorylated S, T, and Y sites, we observed a striking prevalence for these sites to be in low confidence regions (**Figure 5C**), especially when compared to all S, T, and Y residues or even all amino acids. Phosphorylation sites that were detected in all tissues had an even slightly higher preference for these low confidence regions. However, all amino acids observed in non-enriched proteomics experiments did not follow the same trend, emphasizing the specificity of this effect for phosphorylation sites. It is noteworthy that when plotted individually, S, T, and Y phosphorylation sites all exhibit the same trend (**Supplementary Figure 7E and 7F**). These global results confirm previous work that suggests phosphorylation sites are enriched in intrinsically disordered regions.^35,36^ In fact, some efforts have used intrinsically disordered regions to refine phosphorylation site prediction models.^37^ Finally, although phosphorylation is more likely to occur in less structured regions, examination of the surrounding environment shows the confidence scores are increasing with distance from the phosphorylated residue (**Supplementary Figure 7G**). These data suggest further research on the role of structure and phosphorylation site is warranted.

### Kinase predictions based on flanking regions of phosphorylation sites

Further leveraging our extensive dataset of phosphorylation sites across various tissues, we applied a recently published kinase prediction tool^38^, which was developed by using synthetic peptide libraries to profile the substrate sequence specificity of 303 serine/threonine kinases (**Figure 5D**). As expected, a systematic analysis of all phosphorylation sites in our dataset revealed a trend where many phosphorylation sites were assigned to a limited number of putative kinases; however, half of all phosphorylation sites were predicted to be targeted by 24 or more kinases (**Figure 5E**). Furthermore, this approach enabled us to predict kinases for sites that showed tissue enrichment (**Figure 5F**). By comparing enriched phosphorylation motifs between tissue pairs, we identified kinases with established activities in certain tissues (**Supplementary Figure 8**). For instance, comparing brain and liver enriched motifs revealed kinases such as GSK3A^55^, CAMK subfamily 2 members^56–58^, and PCKA/PCKB^59,60^, which are known to modulate the brain phosphoproteome and have been implicated in neurological disease states (**Figure 5G**). While these kinases have activities in other tissues, our dataset emphasized that the phosphorylation sites predicted to be clients of these kinases are strongly overrepresented in brain as compared to liver tissue. Additionally, we identified PHKG2 as the kinase with the strongest frequency factor for liver tissue, consistent with this enzyme’s role as a liver isoform critical for glycogen breakdown.^61^ Our analysis suggests additional kinases with probable tissue-specific activities, highlighting the potential of our resource to facilitate the discovery of such tissue-specific kinase functions.

### Novel phosphorylation sites on mitochondrial proteins

We next sought to investigate how our comprehensive analysis of phosphorylation patterns in the mouse proteome can be further applied in the biochemical and biological context. We first compared our dataset to the mouse phosphorylation dataset from PhosphoSitePlus^52^, revealing a capture of 38,454 previously unidentified phosphorylation sites, constituting 53% of the total identified sites within our dataset (**Figure 6A**). Upon categorizing phosphorylation sites, we observed a significant representation of mitochondrial proteins, with 55% of known mitochondrial proteins harboring at least one phosphorylation site (**Figure 6B**). This observation aligns with the growing appreciation for protein phosphorylation in regulating mitochondrial function.^62^ An exemplary mitochondrial pathway whose protein phosphorylation sites showed the most dynamic pattern across all tissues was the tricarboxylic acid cycle (**Supplementary Figure 9A**) with the majority of sites being detected for the first time in this study (**Supplementary Figure 9B**), setting the stage for further exploration into the understudied regulation of this key metabolic pathway.^63^

**Figure 6.**
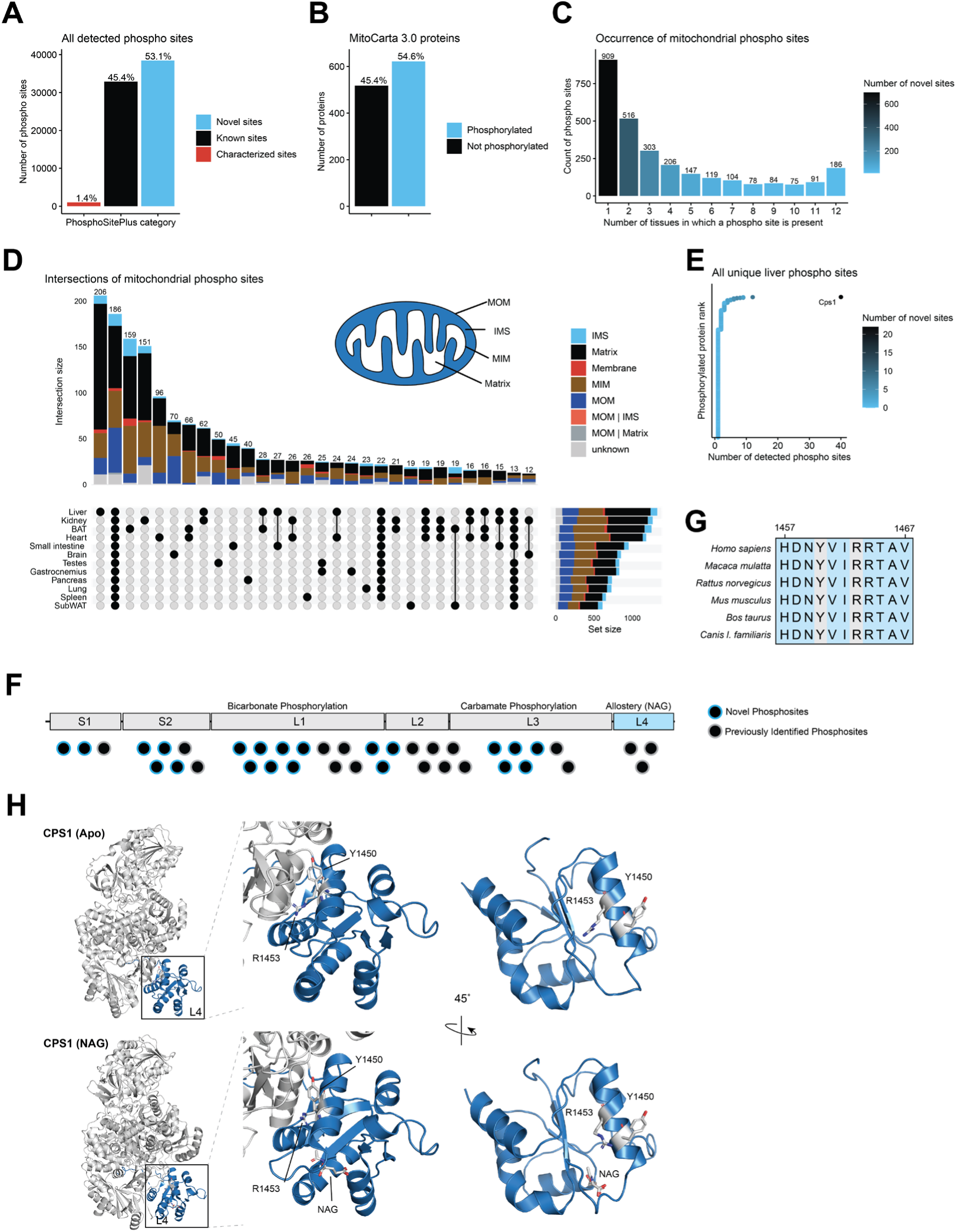
Mitochondrial phosphoproteomics reveals novel liver-specific phosphorylation site. (A) Bar plot of all detected phosphorylation sites in our study stratified into categories that are directly derived from the PhosphoSitePlus database (downloaded August 22, 2023). (B) Mouse MitoCarta 3.0^82^ proteins with at least one detected phosphorylation site in our data versus the remainder. (C) Bar plot indicating the number of mitochondrial phosphorylation sites that occur in a specific number of tissues. (D) Intersections of phosphorylation sites on mitochondrial proteins per tissue subsets. Intersection sizes of 12 or more are shown. Sub-mitochondrial localization is derived from MitoCarta 3.0. (E) Dot plot displaying all proteins that harbor phosphorylation sites unique to liver tissue. The highest ranked protein harbors the most phosphorylation sites as indicated on the x-axis. Novel sites are according to the PhosphoSitePlus database. (F) Schematic representation of the CPS1 peptide chain with all detected phosphorylation sites indicated as novel or previously identified. (G) Sequence alignment of CPS1 orthologues using Clustal Omega^81^. Patient variant residue and phosphorylation site residue of interest are in grey. (H) Structural modeling based on structures (PDB: 5DOT (Apo), 5DOU (NAG)) from RCSB PDB^66^.

The occurrence of phosphorylation sites was variable across tissues in the entire dataset (**Supplementary Figure 9C**) as well as in the mitochondrial subset of our data (**Figure 6C**), indicating that tissue-specific phosphorylation patterns exist within the murine mitochondrial phosphoproteome. To investigate this in more detail, we calculated the intersections of all mitochondrial phosphorylation sites across tissues and found the liver to harbor the most unique phosphorylation sites (**Figure 6D**). Moreover, these unique liver phosphorylation sites showed the largest proportion of mitochondrial matrix proteins, suggesting that phosphoregulation of proteins in this sub-compartment is particularly important in this tissue. We identified carbamoyl phosphate synthetase I (CPS1) to harbor the most phosphorylation sites in this group (**Figure 6E**).

Notably, 40 phosphorylation sites were detected on CPS1, which is the first and rate-limiting enzyme of the urea cycle. None of these phosphorylation sites have been functionally characterized, and 22 of these sites have not been reported previously (**Figure 6F**), underscoring the significant discovery potential of our dataset. Deficiency of human CPS1 results in hyperammonemia ranging from neonatally lethal to environmentally induced adult-onset disease in affected individuals.^64^ The phosphorylation site S913, located within the L2 integrating domain, overlaps with a known severe disease-causing missense variant in CPS1 (p.S913L)^65^, indicating the potential regulatory significance of these sites in CPS1.

Our dataset also provides enhanced phosphotyrosine coverage without specific enrichment methods (**Figure 4B**), revealing novel sites that may be functionally relevant. N-acetyl-L-glutamate (NAG) is an essential allosteric activator of CPS1. By binding to the C-terminal L4 domain of CPS1, NAG induces long-range conformational changes, impacting distant catalytic activities.^66^ We identify Y1450 as a conserved phosphorylated residue in this domain, which is located proximal to the key R1453 residue that is required for NAG binding and is the location of two patient variants (c.4357C>T and c.4358G>A resulting in p.R1453W and p.R1453Q, respectively)^67^ (**Figure 6G**). Previous work suggested that a distinct post-translational modification of Y1450 (nitration) may impact CPS1 activity by obstructing NAG binding.^68^ Structural modeling demonstrates shifts in this Y1450-R1453-containing helix of CPS1 upon NAG binding, highlighting the confirmational change of the R1453 residue and its proximity to the Y1450 residue (**Figure 6H**).

### Most dynamic phosphoregulation in brain tissue

The brain is characterized by a diverse array of expressed kinases and phosphatases and its proteins have a higher tendency to be phosphorylated as compared to other tissues.^49^ Our data substantiates this observation, as brain tissue showed the highest prevalence of unique phosphorylation sites (*i.e.*, sites defined as exclusive to a singular tissue, **Figure 7A**). Moreover, we investigated all pan-tissue expression of phosphorylation sites (sites detected across all twelve tissues) by a z-score analysis, enabling comparison of each phosphosite to the average organismal values, and found brain tissue to exhibit the strongest abundance changes of phosphorylation sites (**Supplementary Figure 10A**). We then applied the same z-score-based principle on the entire dataset (including imputed values for undetected sites in some tissues) to impartially identify sites undergoing significant changes and observed a normal distribution in z-score patterns for all tissues, except the brain, which displayed a pronounced enrichment of phosphorylation sites with elevated z-scores (**Supplementary Figure 10B, Supplementary Table 6**). By applying the same absolute z-score threshold of 2, we found 9,541 phosphorylation site outliers in the brain (**Figure 7B**), accounting for 23% of all outliers across all twelve tissues. Further prioritizing these brain sites based on z-score ranking within the mitochondrial phosphoproteome, we identified serine 298 of Optic atrophy protein 1 (OPA1) as a top hit (**Figure 7C**).

**Figure 7.**
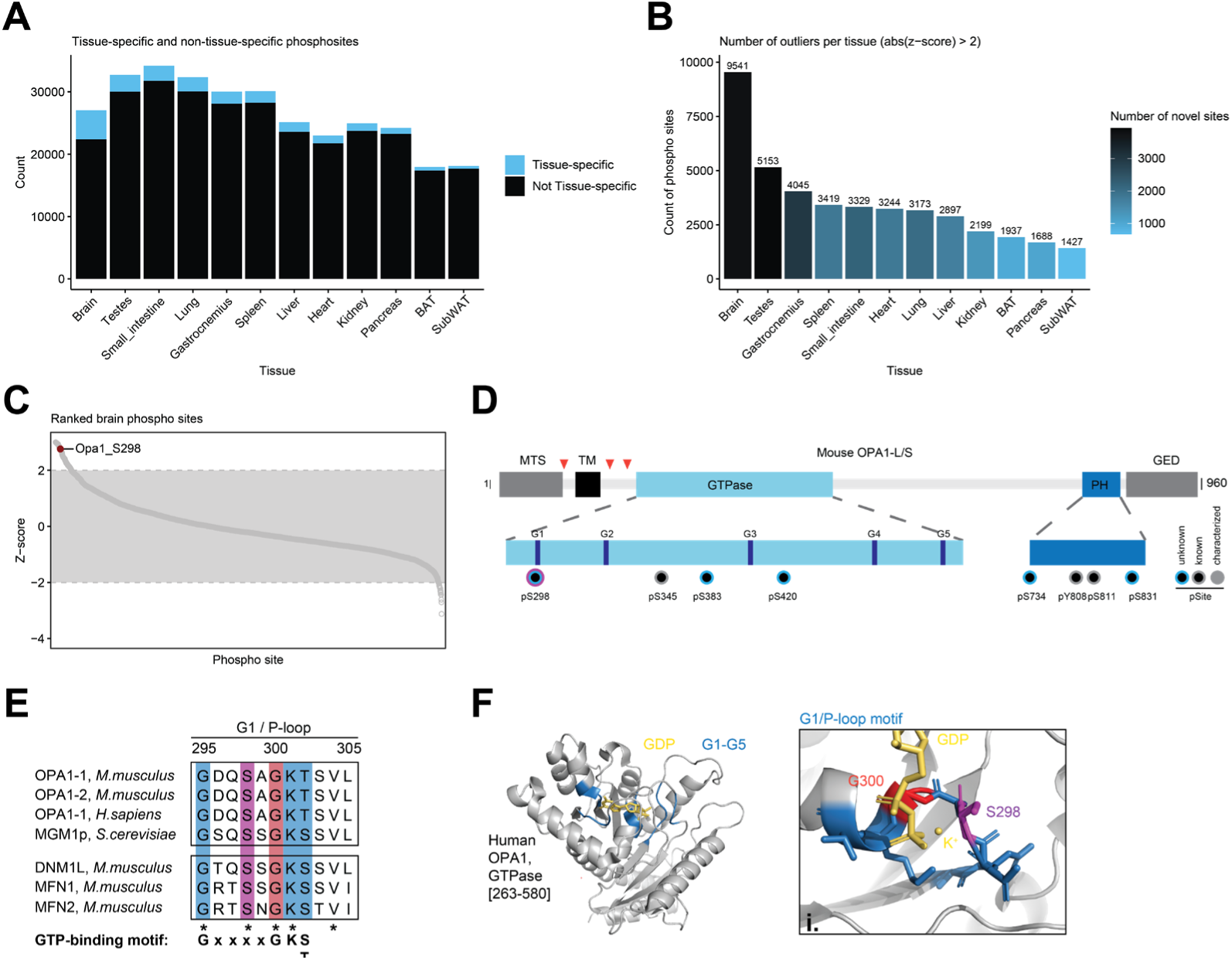
Unique phosphorylation sites in brain tissue. (A) Bar plot indicating all sites (black) that were detected per tissue, highlighting the sites that are unique in each tissue (blue). (B) Number of outliers per tissue based on z-score analysis with an absolute z-score cut-off of 2. (C) Ranked mitochondrial phosphorylation sites in brain based on z-score, highlighting the OPA1 top site. (D) Protein domain representation of mouse OPA1, including mitochondrial transition signal (MTS), transmembrane domain (TM), GTPase, PH, and GED domain. Red triangles represent sites of proteolytic cleavage generating the long and short proteo-forms of OPA1. Note, that all identified phosphosites are common to all OPA1 isoforms. Below, details of GTPase and PH domain with indicated phosphorylation sites (pS/Y) found in brain tissue (See Supplementary Figure 8C for presence of OPA1 phosphorylation sites in other tissues).(E) Sequence alignment of first GTP-binding site/P-loop among mouse OPA1 isoforms and homologs in human and yeast, as well as related dynamin family members, DNM1L, MFN1, and MFN2, in mouse. Conserved residues are marked with *, including serine 298 in mouse OPA1. (F) Structure modeling of human OPA1 GTPase domain [263-580] based on RCSB PDB structure 6JTG^74^. Left, Modeling of the entire domain with highlighted GTP-binding domains (G1-5) in blue and bound GDP (yellow). Right, Detail (i.) of the G1/P-loop domain containing the discussed S298 (magenta) and the nucleotide-binding G300 (red). K+ (yellow) stabilizes the vicinity of the nucleotide.

OPA1 facilitates multiple functions in mitochondria including membrane fusion, cristae biogenesis, mitochondrial DNA maintenance, and respiration.^69^ Importantly, mutations in OPA1 are the most common cause of dominant optic atrophy (DOA). In our data, we detected multiple phosphorylation sites of OPA1 across the twelve tissues (**Supplementary Figure 10C**), none of which are functionally characterized as documented in PhosphoSitePlus.^52^ These sites are localized within the GTPase and PH domain of OPA1, which are also common areas for missense variants in patients with DOA.^70^ Additionally, many of the detected phosphorylation sites have not been previously reported in mouse, among which is the novel S298 site within the first GTP-binding domain (G1/P-loop) of the GTPase (**Figure 7D**). The consensus motif GxxxxGKS/T of G1/P-loop is highly conserved across species and other proteins of the dynamin superfamily (**Figure 7E**). Further, the Q and S residue within this motif are important for the assembly stimulated GTPase activity as shown for dynamin.^71^ Notably, S298 has been identified as a site of missense mutations in DOA patients (c.893G>A, p.S298N^72^; c.892A>G, p.S298G^73^), suggesting its importance for OPA1 function, too. Furthermore, a S298A mutation reduces GTPase activity and dimerization *in vitro*^74^; the S298N mutation abolishes respiratory growth in *S.cerevisiae* with mitochondrial DNA depletion and altered mitochondrial morphology.^75^ Our structural modelling of the OPA1’s GTPase domain shows that S298 is close to the nucleotide-interacting G300 of the G1/P-loop and the nucleotide, suggesting that phosphorylation may affect nucleotide binding or hydrolysis (**Figure 7F**). Taken together, current evidence clearly underscores the functional importance of S298 for OPA1 activity, but the effect and relevance of phosphorylation we have identified on S298 requires further study.

## DISCUSSION

Here we demonstrated that the Orbitrap Astral mass spectrometer provides a platform for fast and deep phosphoproteome analysis. Key to empowering this application is the ability of the system’s high MS/MS duty cycle to enable DIA data collection with narrow isolation windows (2 Th). This capability, combined with high mass accuracy, mass resolution, and sensitive ion detection of the Astral analyzer allow confident phosphosite localization from low abundance peptides as evinced by the performance of the system for low overall phosphopeptide loads. Altogether this approach allowed for the collection of the deepest mouse phosphoproteomic atlas to date following only a half day of data collection.

Nonetheless, the uniquely large mouse phosphoproteome atlas defined here provides an opportunity to explore the sequence and structural context of kinase activity.

From the phosphoproteome analysis we discovered that, despite its bacterial origins, over fifty percent of mitochondrial proteins carry at least one site of phosphorylation site. An interesting and potentially important example is the CPS1 protein on which we detected 22 novel sites. Notably, one of these sites, Y1450, is proximal to the NAG binding domain and near the location of known patient variants – suggesting potential clinical relevance. As exemplified by the above example, this technology permitted detection of thousands of pY sites without the typical pY-specific antibody enrichments. Doubtless the sensitivity and depth afforded by the Orbitrap Astral instrument can allow direct access to these most critical, dynamic, and low-abundance phosphorylation events. Finally, we note that while all tissues exhibit a subset of unique phosphorylation events, the brain is distinguished and contains nearly 10,000 unique phosphorylation sites.

Protein phosphorylation analysis presents many challenges for mass spectrometric analysis; however, we demonstrate here that the Orbitrap Astral resolves many of these limitations and permits fast and deep phosphoproteome analysis. We suppose that these performance characteristics will be translated to the analysis of other post-translational modifications including acetylation, glycosylation, methylation, ubquitinylation, etc. Furthermore, we expect use of extensive fractionation, more tissues, and multiple proteases would allow detection of even more phosphorylation sites and increase tissue-to-tissue phosphorylation site reproducibility.

## ACKNOWLEDGEMENTS

We are grateful for support from the National Institutes of Health (grants P41GM108538 and R35GM118110 to JJC, R01DK098672 to DJP, R35GM150899 to AG, and R35GM147014 to JF), the National Science Foundation grant 2010789, and the Department of Energy grant number DE-SC0018409. PS is supported by a Morgridge Interdisciplinary Postdoctoral Fellowship. TMPC and LRS acknowledge support from the National Human Genome Research Institution through a training grant to the Genomic Science Training Program (NIH T32HG002760). MLR acknowledges support from the UW-Madison Biotechnology Training Program (NIH T32GM135066). TMPC also acknowledges the ACS Division of Analytical Chemistry and Agilent for support through a graduate fellowship. PF acknowledges the support of postdoctoral fellowships by the European Molecular Biology Organization (ALTF 263-2022) and the Swiss National Science Foundation (P500PB_211038). DJP acknowledges the support of funds from the BJC Investigators Program.

## AUTHOR CONTRIBUTIONS

NA and JF performed cell culture and collected HEK293T and HeLa cells. JH and AG collected mouse tissues. JR and APG performed yeast culture and collected yeast cells. TNA, AP, HS, CH, ED, and VZ developed the new instrument used in this study. NML, TMPC, TNA, AP, MLR, LRS, and ED performed mass spectrometry sample preparation and/or data acquisition. NML, PS, TMPC, and LRS performed mass spectrometry data processing. NML, PS, PF, CF, and AJS performed data analysis and assessment of the biological relevance of the study. NML, PS, PF, CF, AJS, ES, TNA, LRS, MSW, ED, VS, and JJC contributed to the figure content and design. NML, PS, PF, CF, AJS, ES, HS, DJP, and JCC wrote the manuscript. NML, PS, PF, TMPC, CF, AJS, ES, TNA, AP, HS, DJP, VZ, and JJC edited the manuscript. MSW, DJP, VZ, and JJC provided project supervision.

## METHODS

### HEK293T Cell Preparation

HEK293T cells (ATCC) were cultured in Dulbecco’s modified Eagle’s medium (DMEM) (Gibco, 11995-065) with 1% penicillin/streptomycin (Thermo Fisher Scientific, 15-140-122) and 10% fetal bovine serum (FBS) (HyClone, 89133-098) at 37°C and 5% CO2. Culture media was replaced every 24 hours. Cells were expanded to appropriate cell number, detached from tissue culture plate with 0.05% trypsin-EDTA (Gibco, 25300062), washed once with phosphate buffered saline (PBS) (Gibco, 02-0119-0500), cell number determined, and 20 x 10^7^ cells pelleted. Cell pellet was immediately stored at –80°C until use. The cells were low passage number and tested negative for mycoplasma contamination. Frozen cell pellets were resuspended in 5.4M guanidine hydrocholoride (from Sigma Life Science, 8M, pH 8.5, G7294-100mL) in 100 mM Tris, pH 8 (Invitrogen, 1 M Tris pH 8.0, 0.2 µm filtered, AM9856) via vortexing, followed by heating in a sand bath for 5 minutes at 105℃ prior to brief (10-15s) sonication with a probe sonicator. The sample was diluted with the guanidine buffer above to give a ∼1.5 mg/mL estimated protein concentration via NanoDrop (Thermo Scientific) prior to beginning digestion.

### EGF-stimulated HeLa Cell Preparation

HeLa cells were cultured in Dulbecco’s Modified Essential Medium (DMEM) (Gibco, 11995-065) with 10% FBS (HyClone, 89133-098) and 1% penicillin/streptomycin (Thermo Fisher Scientific, 15-140-122) in 37°C/5% CO2 incubator. Cells were resuspended in PBS (Gibco, 02-0119-0500) with or without 100 ng/mL human Endothelial Growth Factor (Thermo, AF-100-15) for 15 minutes at room temperature. Cells were gently resuspended every 5 minutes during incubation. After incubation, cells were washed twice with PBS and stored in –80°C until use. Protein extraction was performed as described above for the HEK293T samples. Protein concentrations were estimated via protein BCA (Pierce, 23235).

### Yeast Cell Preparation

*Saccharomyces cerevisiae S288C-derivative strain, BY4741,* was cultured in triplicate for ∼5 generations into log phase in rich YPD medium at 30°C. Cells were centrifuged at 3000 RPM, 3 min, rinse in water and snap-freeze in liquid nitrogen. Frozen cell pellets were resuspended in 8M urea Sigma-Aldrich, U5378) in 100 mM Tris, pH 8 (Invitrogen, 1 M Tris pH 8.0, 0.2 µm filtered, AM9856) and vortexed with glass beads (425-600 µm, Sigma-Aldrich, G8772-500G) to lyse (2 minutes of total vortexing with 30s vortexing followed by 30s on ice). Protein concentrations were estimated via protein BCA (Pierce, 23235).

### Mouse Tissue Preparation

All experiments were performed in accordance with the National Institute of Health Guide for the Care and Use of Laboratory Animals and were approved by the Animal Care and Use Committee at the University of Wisconsin-Madison. Six-week-old male C57BL6/J mice (n=3) were euthanized by cervical dislocation and tissues were immediately collected, and flash frozen in liquid nitrogen. A total of twelve tissues (pancreas, small intestine, spleen, liver, kidney, testes, heart, lung, subcutaneous white adipose tissue (WAT), brown adipose tissue (BAT), gastrocnemius, and brain) were collected. All tissues were stored at –80°C prior to cryo-pulverization. For each tissue, samples from three mice were pulverized together into a fine power under liquid nitrogen. For each pulverized tissue, ∼60mg frozen wet weight was resuspended in 4 mL of 5.4M guanidine hydrocholoride (Sigma Life Science, 8M, pH 8.5, G7294-100mL) in 100 mM Tris, pH 8 (Invitrogen, 1 M Tris pH 8.0, 0.2 µm filtered, AM9856) with Pierce™ Phosphatase Inhibitor Mini Tablets (A32957) with one tablet/10mL. Tissue samples were vortexed and sonicated for 20 min in a bath sonicator (chilled to 4°C) to homogenize. A probe sonicator was used briefly to sonicate samples on ice as needed. The protein concentration was estimated via protein BCA (Pierce, 23235) and samples were diluted with the guanidine hydrochloride buffer above to give a protein concentration of ∼2mg/mL prior to digestion.

### Protein digestion

After extracting proteins from human cells, yeast cells, or mouse tissues, methanol (Optima LC/MS grade, Fisher Scientific) was added to 90% (v/v) to precipitate protein and samples were vortexed prior to centrifugation at 4000g for 15 minutes. The supernatant was removed, and the pellet was resuspended in 8M urea (Sigma-Aldrich, U5378), 100mM Tris (Invitrogen, 1 M Tris pH 8.0, 0.2 µm filtered, AM9856), 10mM TCEP (Sigma-Aldrich, C4706-2G), 40mM 2-chloroacetamide (Sigma Aldrich, ≥98%, C0267-100G) pH 8 at ∼1.5mg protein/mL. Lysyl Endopeptidase (LysC, 100369-826, VWR) was added at a ratio of 1:50 enzyme:protein and gently rocked at ambient temperature for 4 hours, followed by the dilution of the solution to 2M urea with 100mM Tris, pH 8 (Invitrogen, 1 M Tris pH 8.0, 0.2 µm filtered, AM9856). Promega Sequencing Grade Modified Trypsin (V5113) was added at a ratio of 1:50 enzyme:protein and incubated overnight. Following overnight digestion, the solution was acidified to <pH 2 with 10% trifluoroacetic acid (Sigma-Aldrich, HPLC grade, >99.9%) to quench the digestion. The sample was then centrifuged at 4000g for 10 minutes to remove particulate matter prior to desalting with a Strata-X 33 µm polymeric reversed phase SPE cartridge. Peptides were dried via a SpeedVac (Thermo Scientific) and stored at –80°C until phosphopeptide enrichment or, in the case of the mouse proteomics experiments, until fractionation.

### Phosphopeptide enrichment

Phosphopeptides were enriched from digested peptides using MagReSyn Ti-IMAC HP beads (ReSyn Biosciences, MR-THP005). A volume of 100µL beads were used per 1mg of peptides. Input peptide masses of 2-3mg were utilized for human and yeast enrichments, and input peptide masses ranging from ∼0.3 mg to 1.2 mg were utilized for the different mouse tissues. Beads were washed three times with 1 mL 80% acetonitrile (Optima LC-MS grade, Fisher Scientific)/6% trifluoracetic acid (Sigma-Aldrich, HPLC grade, >99.9%) prior to resuspending the sample in 1 mL 80% acetonitrile (Optima LC-MS grade, Fisher Scientific)/6% trifluoracetic acid (Sigma-Aldrich, HPLC grade, >99.9%) and vortexing the sample with the beads for 1 hour. After the 1-hour of vortexing, the beads were washed three times with 1mL 80% acetonitrile (Optima LC-MS grade, Fisher Scientific) /6% trifluoracetic acid (Sigma-Aldrich, HPLC grade, >99.9%), once with 1 mL 80% acetonitrile, once with 1 mL 80% acetonitrile (Optima LC-MS grade, Fisher Scientific) /0.5 M glycolic acid (Sigma-Aldrich, 99%, 124737-500G), and three times with 1 mL 80% acetonitrile (Optima LC-MS grade, Fisher Scientific). The phosphopeptides were eluted from the beads with the addition of 300 µL 50% acetonitrile (Optima LC-MS grade, Fisher Scientific)/1% ammonium hydroxide (28% in H2O, ≥99.99% trace metals basis, Sigma-Aldrich), followed by a second elution with another 300 µL 50% acetonitrile/1% ammonium hydroxide. The samples were acidified via addition of 15 µL 10% trifluoroacetic acid (Sigma-Aldrich, HPLC grade, >99.9%). The samples were then dried down in a SpeedVac (Thermo Scientific) prior to being resuspended in 0.2% trifluoroacetic acid (Sigma-Aldrich, HPLC grade, >99.9%) and desalted as previously described. Desalted phosphopeptide samples were dried and resuspended in 0.1% formic acid (Fisher Scientific, LC-MS grade). The phosphopeptide concentration was estimated via NanoDrop (Thermo Scientific). The HEK293T phosphopeptides were pooled to generate a sample for method evaluation. Each mouse phosphopeptide sample was fractioned as described below.

### High-pH Peptide Fractionation

High-pH fractionation of peptides was performed on an Agilent 1260 Infinity BioInert LC with an automated fraction collector. A 20-minute method was performed on a Waters XBridge, Peptide BEH C18, 3.5 µm, 130 Å, 4.6mm x 150 mm column with a flow rate of 0.8 mL/min. Mobile phase A and B were 10 mM ammonium formate (Sigma-Aldrich, >99/0%, LC-MS grade, 70221-100GF), pH 10 and 20% 10mM ammonium formate (pH 10)/80% methanol (Optima LC/MS grade, Fisher Scientific), respectively. The gradient went from 0 to 35%B from 0-2 minutes, 35 to 75%B from 2-8 minutes, 75% to 100%B from 8-13 minutes, followed by washing at 100%B from 13-15 minutes and equilibration at 0%B from 15-20 minutes. UV absorbance at 210 and 280nm was recorded. For HEK293T spectral library generation and mouse proteomics samples, 16 fractions were collected from 5 to 18 minutes and concatenated into the final fractions by combining fraction 1 and 9, fraction 2 and 10, etc., resulting in 8 final fractions. For mouse phosphoproteomics samples, 8 fractions were collected from 5 to 18 minutes and concatenated into the final fractions by combining fraction 1 and 5, fraction 2 and 6, etc. Samples were dried down in a SpeedVac (Thermo Scientific) prior to being resuspended in 0.1% formic acid (Fisher Scientific, LC-MS grade) for LC-MS analysis.

### Synthetic Phosphopeptide Dilution Series Preparation

Five sets of phosphopeptide standards were acquired: SpikeMix PTM-Kit 52 (JPT, SPT-PTM-POOL-Yphospho-1), SpikeMix PTM-Kit 54 (JPT, SPT-PTM-POOL-STphospho-1), MS PhosphoMix 1 (Sigma, MSP1L-1VL), MS PhosphoMix 2 (Sigma, MSP2L-1VL), and MS PhosphoMix 3 (Sigma, MSP3L-1VL). The standards were reconstituted in 0.2% formic acid/20% acetonitrile/80% water via vortexing. The standards were pooled into an equimolar mixture of the 225 total phosphopeptide standards. The pooled equimolar mixture was then diluted and mixed with the yeast phosphopeptide sample to construct a dilution series comprised of five points of four-fold dilutions starting at 10000 amol, resulting in total quantities loaded onto the column of 10000, 2500, 625, 156.25, and 39.0625 amol per phosphopeptide standard along with a constant yeast phosphopeptide load of 250ng. A summary of all the phosphopeptide standards used in the analysis is provided in **Supplementary Table 1**.

### LC-MS Operation for Phosphoproteomics with Orbitrap Astral Analysis

Nanoflow capillary columns (75µm I.D., 360µm O.D.) with pulled nanoESI emitters were packed to 40cm at high pressures with C18 1.7 µm diameter, 130 Å pore size BEH C18 particles (Waters) as previously described.^39^ Samples were analyzed with a Vanquish Neo UHPLC coupled to an Orbitrap Astral mass spectrometer (Thermo Scientific) using a NanoSpray Flex source (Thermo Scientific). A source voltage of 2000 V was used for all experiments. Mobile phase A and B were 0.1% formic in water (Fisher Scientific, Optima LC-MS grade) and 0.1% formic acid/80% acetonitrile (Fisher Scientific, Optima LC-MS grade), respectively. The column was heated to 50°C with the Column Oven PRSO-V2 (Sonation Lab Solutions) and the flow rate was set to 400nL/min at the start of the method to decrease delay time and turned to 300nL/min at the start of the active gradient. Initial conditions of 2%B were ramped to 14% from 0 to 5 minutes. The active gradient was generally set to 14% to 54%B with curve type 6 beginning at 5.2 minutes, with the exact %B settings adjusted for each active gradient length to evenly distribute peptide signal across the gradient. The column was washed for 5 minutes at 100%B and 400 nL/min at the end of the gradient, followed by fast equilibration on the Vanquish Neo LC with an upper pressure limit of 1100 bar.

For initial DDA experiments on the Orbitrap Astral MS, MS^1^ spectra were collected in the Orbitrap every 0.6s at a resolving power of 240,000 at m/z 200 over *m/z* 350-1350 with a normalized AGC target of 300% (3e6 charges) and a maximum injection time of 10ms. The MIPS filter was applied with Peptide mode and ‘Relax Restrictions when too few Precursors are Found’ set to True. Precursors were filtered to charges states 2-6. A Dynamic Exclusion filter was applied with 10s duration and 10ppm low and high mass tolerance and exclude isotopes set to True. An intensity filter was applied with a minimum precursor intensity of 5000 required for selection. MS^2^ scans were collected in the Astral mass analyzer with an isolation window of 0.7 *m/z*, normalized collision energy of 27, a scan range of 150-2000 *m/z*, an AGC target of 100% (1e4 charges), and a maximum injection time of 10ms.

For DIA experiments on the Orbitrap Astral MS, MS^1^ spectra were collected in the Orbitrap every 0.6s at a resolving power of 240,000 at m/z 200 over *m/z* 380-980. The MS^1^ normalized AGC target was set to 300% (3e6 charges) with a maximum injection time of 10ms. DIA MS^2^ scans were acquired in the Astral analyzer over a 380-980 m/z range with a normalized AGC target of 500% (5e4 charges) and a maximum injection time of 3.5ms and an HCD collision energy setting of 27% and a default charge state of +2. Window placement optimization was turned on. Isolation widths of 2 Th and active gradient lengths of 30 minutes were used with HEK293T fraction analysis for HEK293T spectral library generation. Isolation bin widths of 2 Th and 4 Th with 1 Th overlap were compared for HEK293T phosphoproteomics method evaluation. Isolation widths of 2 Th were used for mouse phosphoproteomics experiments.

### LC-MS Operation for Benchmarking on the Orbitrap Ascend

Nanoflow capillary columns (75µm I.D., 360µm O.D.) with pulled nanoESI emitters were packed to 40cm at high pressures with C18 1.7 µm diameter, 130 Å pore size BEH C18 particles (Waters) as previously described.^39^ Samples were analyzed with a Vanquish Neo UHPLC (Thermo Scientific) coupled to an Orbitrap Ascend mass spectrometer (Thermo Scientific) using a NanoSpray Flex source (Thermo Scientific) incorporating a homebuilt column heating compartment. The column was heated to 50°C. The flow rate was set to 400nL/min at the start of the method to decrease delay time and turned to 300nL/min at the start of the active gradient. Initial conditions of 2%B were ramped to 14% from 0 to 5 minutes. The active gradient was generally set to 14% to 54%B with curve type 6 beginning at 5.2 minutes, with the exact %B settings adjusted for each active gradient length to evenly distribute peptide signal across the gradient. The column was washed for 5 minutes at 100%B and 400 nL/min at the end of the gradient, followed by fast equilibration on the Vanquish Neo LC with an upper pressure limit of 1100 bar.

DIA experiments on the Orbitrap Ascend utilized similar method settings to those previously described by Bekker-Jensen et al^41^, but with the same *m/z* range as utilized on the Orbitrap Astral to enable direct comparisons. Briefly, MS^1^ spectra were collected in the Orbitrap at a resolving power of 60,000 at m/z 200 over *m/z* 380-980 with a AGC target of 250% and maximum injection time of 123 ms. DIA MS^2^ scans were collected in the Orbitrap with 15,000 at 200 *m/z* resolving power, a scan range of 150-2000 *m/z*, an AGC target of 200%, a maximum injection time of 27 ms, and a collision energy of 30%. A *m/z* range from 380-980 *m/z* was iterated through with 14 Th isolation windows with 1 Th overlap.

### LC-MS Operation for Proteomics with Orbitrap Eclipse Analysis

The chromatography setup described above for the Orbitrap Ascend analysis was utilized for the Orbitrap Eclipse experiments. A source voltage of 2000 V was used for all experiments. Mobile phase A and B were 0.2% formic acid (Optima LC-MS grade, Fisher Scientific) in water (Optima LC-MS grade, Fisher Scientific) and 0.2% formic acid/80% acetonitrile (Optima LC-MS grade, Fisher Scientific), respectively. The column was heated to 50°C and a flow rate of 300 nL/min was used. Initial conditions of 0%B were ramped to 6%B from 0 to 1 minute. The active gradient was set to 6% to 52%B with curve type 6 from 1 to 73 minutes. The column was washed for 5 minutes at 100%B, followed by fast equilibration on the Vanquish Neo LC with an upper pressure limit of 1150 bar.

For DDA experiments on the Orbitrap Eclipse MS, MS^1^ spectra were collected in the Orbitrap analyzer every 0.6s at a resolving power of 240,000 at m/z 200 over *m/z* 300-1350 with a normalized AGC target of 250% and a maximum injection time of 50ms. The MIPS filter was applied with Peptide mode and ‘Relax Restrictions when too few Precursors are Found’ set to True. Precursors were filtered to charges states 2-5. A Dynamic Exclusion filter was applied with 10s duration and 25ppm low and high mass tolerance and exclude isotopes set to True. MS^2^ scans were collected in the ion trap with an isolation window of 0.5 *m/z*, normalized collision energy of 25, a scan range of 150-1350 *m/z*, an AGC target of 300%, and a maximum injection time of 14ms.

### DDA Data Processing

Proteomics DDA data was processed in MaxQuant 2.4.9.0 with default parameters.^40^ Phosphoproteomics DDA data was processed with the same version of MaxQuant with default parameters and Phospho (STY) enabled as a variable modification. The ‘Phospho (STY)Site.txt’ was used for analysis with a filter for localization probabilities greater than or equal to 0.75.

### DIA Data Processing

Phosphoproteomics DIA data was processed in Spectronaut version 17.6.230428.55965 or 18.6.231227. A HEK293T spectral library was generated in Spectronaut (searching against the human proteome database (Swiss-Prot and TrEMBL) downloaded from UniProt on 15-Jan-2023) from the eight HEK293T phosphopeptide fractions analyzed with a 30-minute active gradient 2 Th DIA method and the library was used to search the HEK293T phosphoproteomics method evaluation experiments shown in Figure 2. Note that the entrapment search in Figure 2G and the phosphoproline variable modification search in Figure 2H were performed as a library-free search. Mouse phosphoproteomics data was searched with the directDIA mode. Default Spectronaut search parameters were used with a variable Phospho (STY) modification included. The PTMSiteReport file was used for analysis of phosphorylation sites. To count the number of unique phosphorylation sites detected within an experiment, the PTMSiteReport was filtered for rows with ‘Phospho (STY)’ in the ‘PTM.ModificationTitle column and values greater than or equal to 0.75 in the ‘PTM.SiteProbability’ column. The data table was then sorted according to ‘PTM.CollapseKey’ and filtered for unique values in the ‘PTM.Group’. At this stage, there can be rows with the same values of ‘PTM.CollapseKey’ that are not filtered by ‘PTM.Group’ due to PTM grouping differing across multiple files, so the dataset was filtered for unique values of ‘PTM.CollapseKey’ to determine the total number of unique phosphorylation sites measured in a single file. Mouse phosphoproteomics data was searched against the mouse proteome downloaded from UniProt from Swiss-Prot and TrEMBL (downloaded on 25-Aug-2023). For the mitochondrial biology investigation, a separate search was performed with just the Swiss-Prot database to facilitate the analysis.

HEK293T phosphoproteomics DIA data was also processed in Proteome Discoverer 3.1.0.638 with CHIMERYS. Numbers of localized phosphosites were reported from the Modification Sites table after filtering for rows with ‘Phospho’ in the Modification Name column. These searches were performed in a developmental version of Proteome Discoverer 3.1 to show that the phosphoproteomic depth achieved with our methods is not restricted to Spectronaut searches. However, this developmental version does not yet incorporate quantitative values for phosphorylation sites detected with DIA methods. Furthermore, from internal conversations, we know the results of this development build should be treated as preliminary and that the algorithms being utilized are still in active development. Consequently, we chose to perform our analyses using Spectronaut, which provides identification, localization, and quantification for DIA phosphoproteomics.

### Localization Error Rate Calculation

To calculate the localization error rate, a reference sheet of all the possible precursors that could be detected from the phosphopeptide standards was constructed with the phosphorylation sites specified. Precursors with sequences that could also arise from the yeast proteome (based on an *in silico* digest) were removed from consideration.

Then, the error rate was calculated as Error Rate 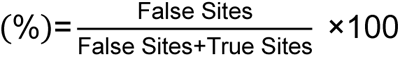, where ‘False Sites’ is the number of phosphopeptide standard precursors detected with phosphorylation states not indicated in the reference sheet, and ‘True Sites’ is the number of phosphopeptide standard precursors detected with phosphorylation states indicated in the reference sheet. This calculation was performed for each phosphopeptide standard raw data file, and the average error rates are reported as a function of the localization probability cutoff.

### Phosphopeptide Standard Quantification Calculations

The phosphopeptide standard reference sheet described above was modified so that each row corresponds to a phosphorylation site, so that the ‘PTM.Quantity’ value in the PTM Site Report could be utilized for quantification assessment. As described above, phosphopeptides with sequences that could arise from the yeast proteome were removed from consideration. Linear regression and R^2^ calculations were performed using the ‘pearsonr’ and ‘linregress’ functions within the ‘scipy.stats’ module for phosphosites detected across at least three concentration points.^76^

### EGF-stimulated HeLa Differential Expression and Pathway Analysis

Differential expression analysis for the EGF treatment experiment was performed on all phosphosites that were detected across triplicates in either the EGF-treatment or control group. Missing value imputation was performed in Perseus^77^ assuming a normal distribution with ‘width = 0.3’ and ‘down shift = 1.8’. Enriched phosphosites were defined as those with an absolute fold change of more than 2 and a p-value less than 0.05.

Pathway enrichment analysis was performed using Enrichr.^78^ The enriched subset contains genes corresponding to phosphosites that are enriched as defined above, and the background was performed for the rest of genes with at least one detected phosphosite.

### Sequence-based phosphosite clustering

Phosphosite sequences, along with their 5-amino acid flanking regions, were compared using the Blosum62 substitution matrix. The top 4 scores, representing a typical number of amino acids in kinase recognition motifs, were selected. These scores were subsequently averaged and transformed to a range between 0 (indicating identical sequences) and 1 (representing maximally different sequences). Subsequently, a similarity matrix between all sequence pairs was constructed and utilized to generate a 2D embedding space through t-SNE, implemented using the scikit-learn library.^79^ Default parameters were used, except for metric=“precomputed”, init=“random”, n_iter=500, n_iter_without_progress=150, and random_state=42. Additionally, Perseus was employed for visualization purposes.^77^

### Kinase Motif Enrichment

Initially, each phosphorylation site and its flanking region in our dataset was used to search the kinase library website (https://kinase-library.phosphosite.org/site, accessed on February 22nd, 2024) for predictions of all 303 serine/threonine kinases for each site. For kinase motif enrichment analysis, only the top 15 kinases with regards to percentile score for each site were retained. To nominate tissue-enriched phosphorylation sites, we proceeded as follows: For each tissue-tissue comparison the log-transformed abundances of all shared phosphorylation sites were subtracted from each other. For selection of top phosphorylation sites per tissue two different approaches were applied: a z-score threshold of 2 was used, or the top and bottom 200 sites were selected. For the final analysis, the top and bottom 200 ranked sites were used. To calculate the frequency factor for each kinase the proportion of how often a kinase was predicted within the top phosphorylation sites versus the total of top phosphorylation sites was computed, divided by the same ratio for the unchanged sites, i.e., not ranked among the top/bottom 200. A chi-squared test was calculated after applying Haldane’s correction to the contingency table of the two proportions and the p-value was extracted.

### Human variant information

Annotation of human variants in CPS1 and OPA1 were identified using The Human Gene Mutation Database at the Institute of Medical Genetics in Cardiff (HGMD) using the public site entries (retrieval date 2023-11-18 ^80^).

### Multiple Sequence Alignment

Clustal Omega^81^ hosted at the EMBL-EBI webserver was used for protein sequence alignment using the following UniProt identifiers for CPS1: P08686, F7EZ, P07756, Q8C196, F1ML89, A0A8C0NKH0; for OPA1: P58281, P58281-2, O60313, P32266; and other dynamin family members: Q811U4, Q80U63, Q8K1M6 (retrieved from UniProt 2023-11-18).

### Structural modelling prediction

The PyMOL Molecular Graphics System, Version 2.5.7 Schrödinger, LLC. was used for predicting structural models for CPS1 (RCSB PDB: 5DOT (Apo), 5DOU (NAG)^66^) and OPA1 (RCSB PDB structure 6JTG ^74^). For modeling purposes, the human crystal structures (highly homologous to mouse) were used.

### Code Availability

All code will be available at the time of publication.

### Data Availability

All data will be available at the time of publication.

## SUPPLEMENTARY INFORMATION

**Supplementary Figure 1.**
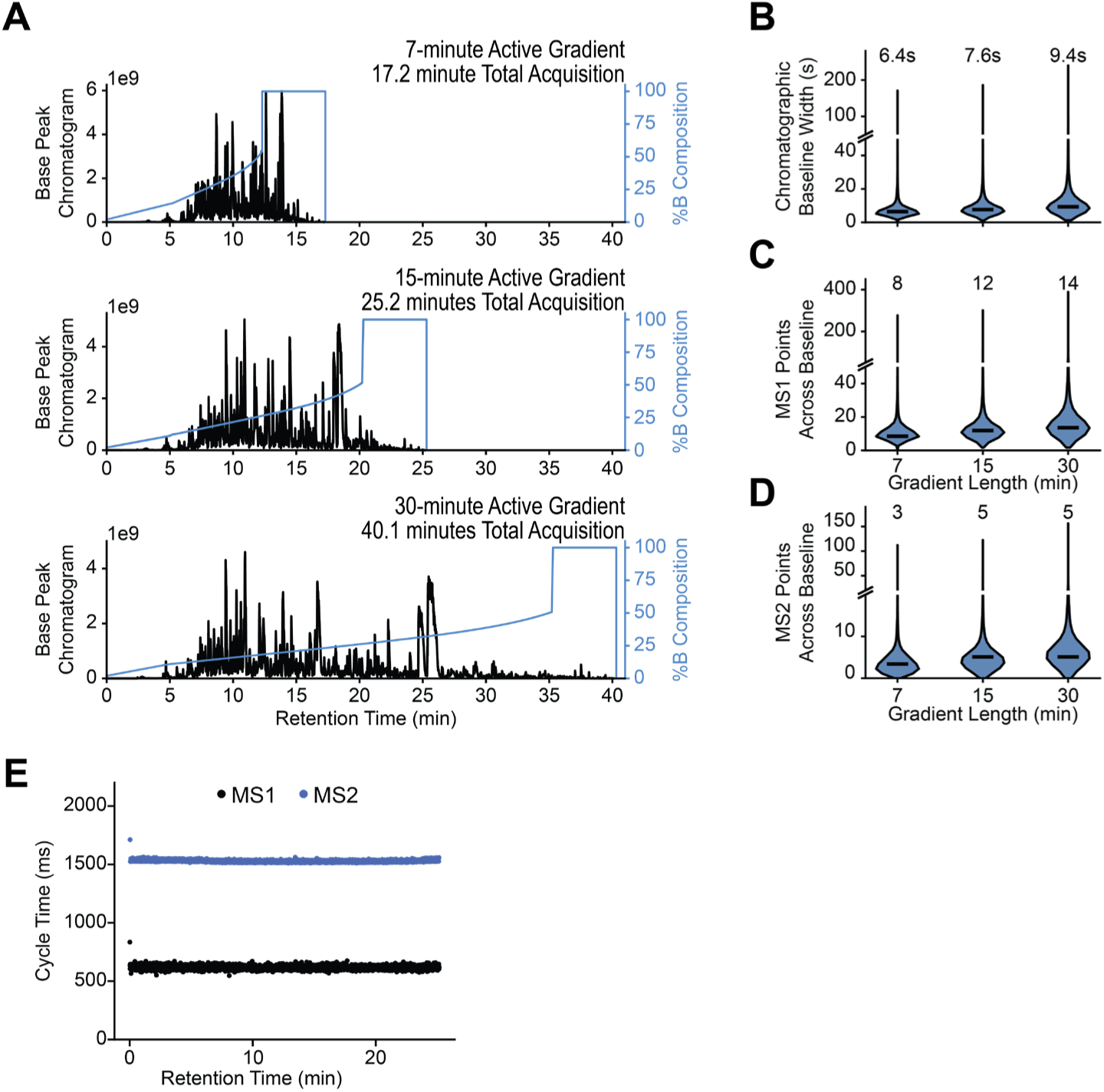
Chromatographic Properties of HEK293T Phosphoproteomics Datasets. (A) Representative chromatograms for different gradient lengths. The base peak chromatograms for the three different active gradient lengths are shown with the gradient composition curve plotted. The distribution of (B) chromatographic baseline widths, (C) MS1 points across the baseline peak, and (D) MS2 points across the baseline peaks are shown with the median value indicated in the violin plot and printed above for three different active gradient lengths. The values were estimated by multiplying chromatographic FWHM, MS1 points across FWHM, and MS2 points across FWHM reported by Spectronaut 17 by 1.7 (based on the 4σ baseline width of a Gaussian distribution where 4σ ≈1.7 * FWHM). (E) The cycle times for the MS1 and MS2 scan sequences are shown as a function of RT.

**Supplementary Figure 2.**
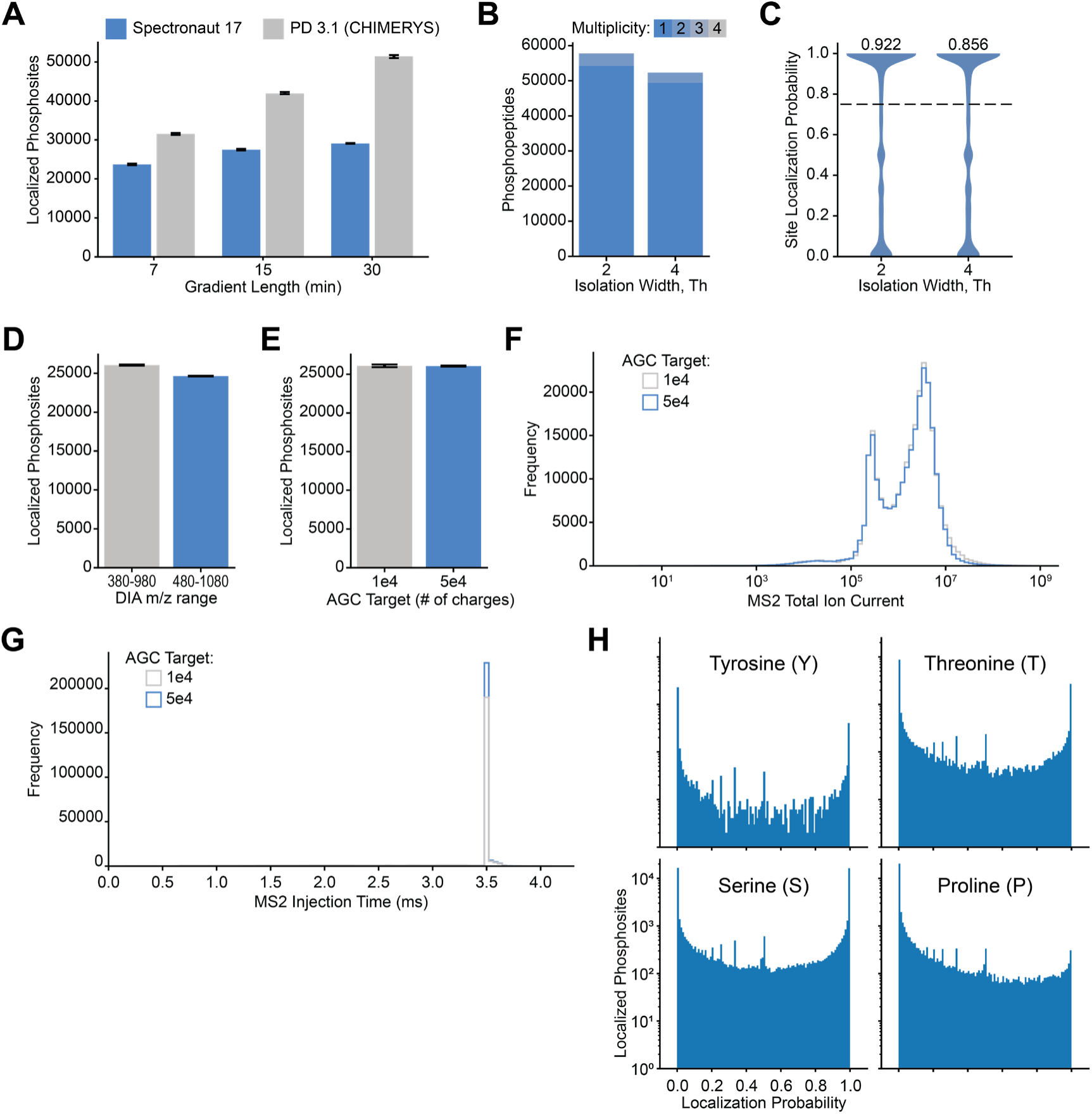
Evaluation of Phosphoproteomic Method and Processing Parameters. (A) The average number of localized phosphosites across triplicate injections for three different gradient lengths is shown for searches in Spectronaut (blue) and CHIMERYS in Proteome Discoverer 3.1 (gray). The error bars represent the maximum and minimum value across the triplicates. (B) The number of phosphopeptides and the multiplicity distributions are shown for the data in Figure 2B. (C) The site localization probability distributions are shown for the data in Figure 2B with the median value indicated above. (D) The average number of phosphosites detected across two replicate injections in EGF-stimulated HeLa cells are shown for two different DIA *m/z* ranges. Error bars indicated the maximum and minimum number of phosphosites. (E) The average number of phosphosites detected across two replicate injections in EGF-stimulated HeLa cells are shown for two different AGC target values. Error bars indicated the maximum and minimum number of phosphosites. (F) The MS2 total ion current distributions are indicated for the two AGC targets in **Supplementary Figure 2E**. (G) The distributions of MS2 injection times are indicated for the two AGC targets in **Supplementary Figure 2E**. (H) Histograms of localization probabilities for S, T, Y, and P residues are shown as an alternative visualization for the phosphoproline decoy search results in Figure 2H.

**Supplementary Figure 3.**
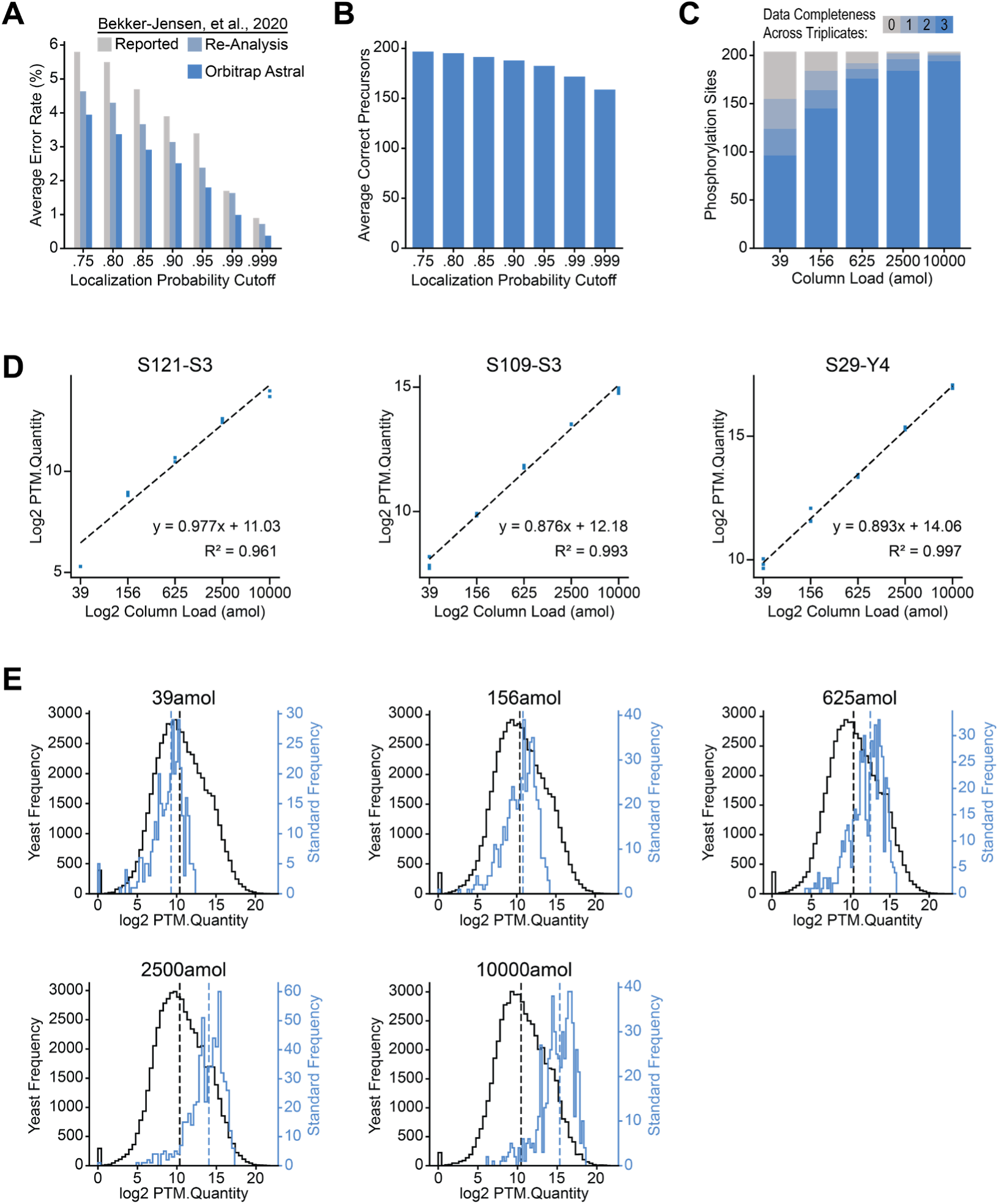
Validation of Site Localization and Quantification using Synthetic Phosphopeptide Standards. (A) The localization error rates shown in Figure 2I are compared to the values previously reported by Bekker-Jensen *et al*^41^ alongside the error rates calculated following re-analysis of the data from Bekker-Jensen *et al* with the same protein database and Spectronaut workflow used in this study. (B) The average number of correctly identified precursors as a function of localization probability cutoff are shown for the dataset in Figure 2I. (C) The detection completeness of synthetic phosphosites across triplicate injections is shown as a function of the column load for the data in Figure 2J. (D) Calibrations curves for three randomly selected synthetic phosphosites are shown from the synthetic phosphopeptides dilution series dataset. ‘S121-S3’ indicates the phosphorylation serine at position 3 for phosphopeptide standard 121. (E) The distribution of phosphosite intensities (Spectronaut’s PTM.Quantity value) for the synthetic phosphopeptide standards and yeast phosphopeptides are shown in blue and black, respectively, for each of the dilution series points. The median intensity for the phosphopeptide standards and yeast phosphopeptides is indicated on each plot with a blue and black dotted line, respectively.

**Supplementary Figure 4.**
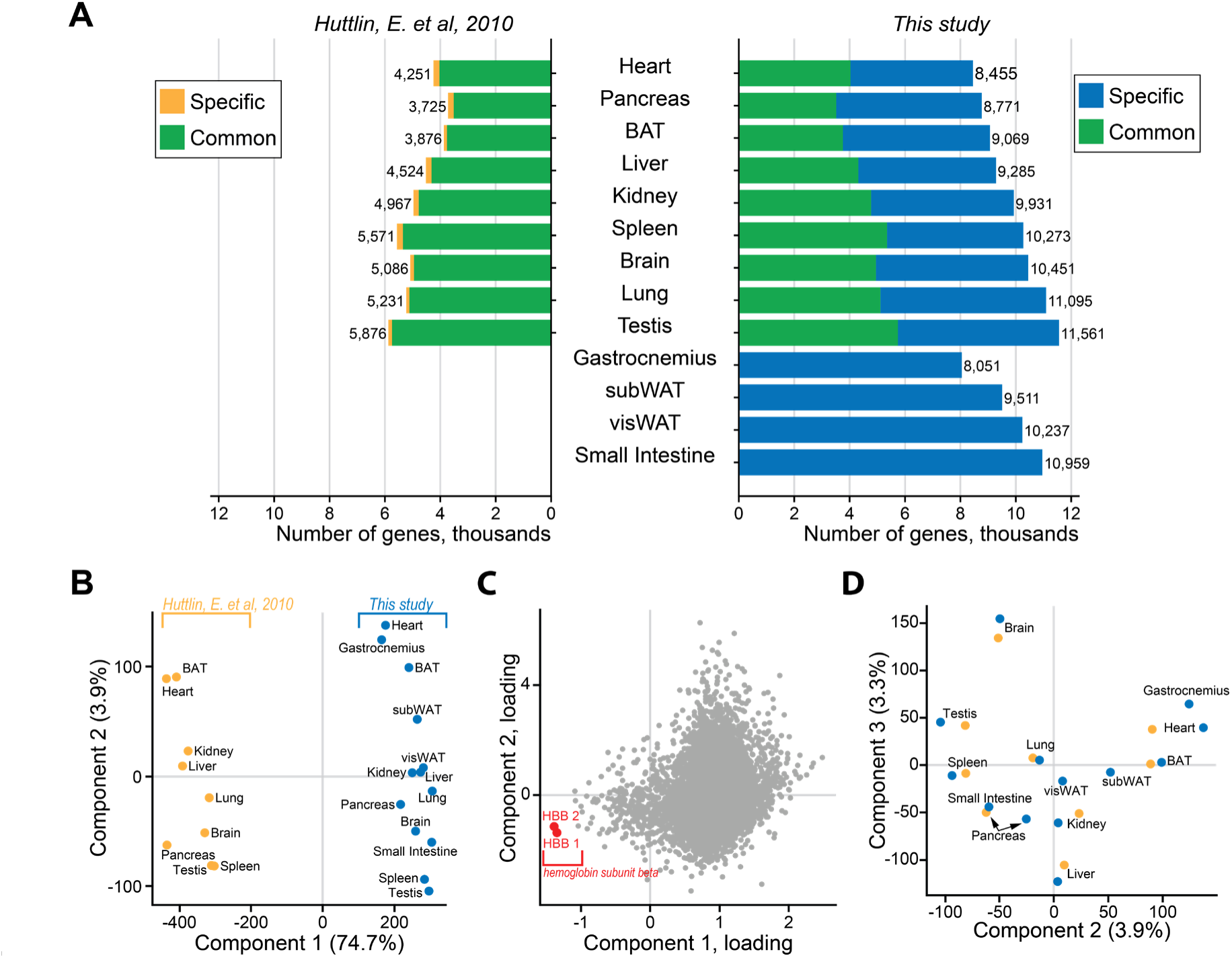
Proteomic Profiling of Mouse Tissues. (A) Results from Mouse Tissue Proteomic Analysis. The numbers of protein groups are indicated for each tissue, with common protein groups highlighted in green, those unique to Huttlin et al. in yellow, and those unique to this study in blue. The Huttlin et al. results were generated by analyzing raw data in MaxQuant, employing the same protein database utilized in this study. (B) Principal Component Analysis (PCA) based on protein expression of common proteins between two studies. (C) PCA loading plot of first two components from (B) with highlights of HBB1 and HBB2 proteins. (D) The same PCA as (B) but Component 2 and Component 3 visualized.

**Supplementary Figure 5.**
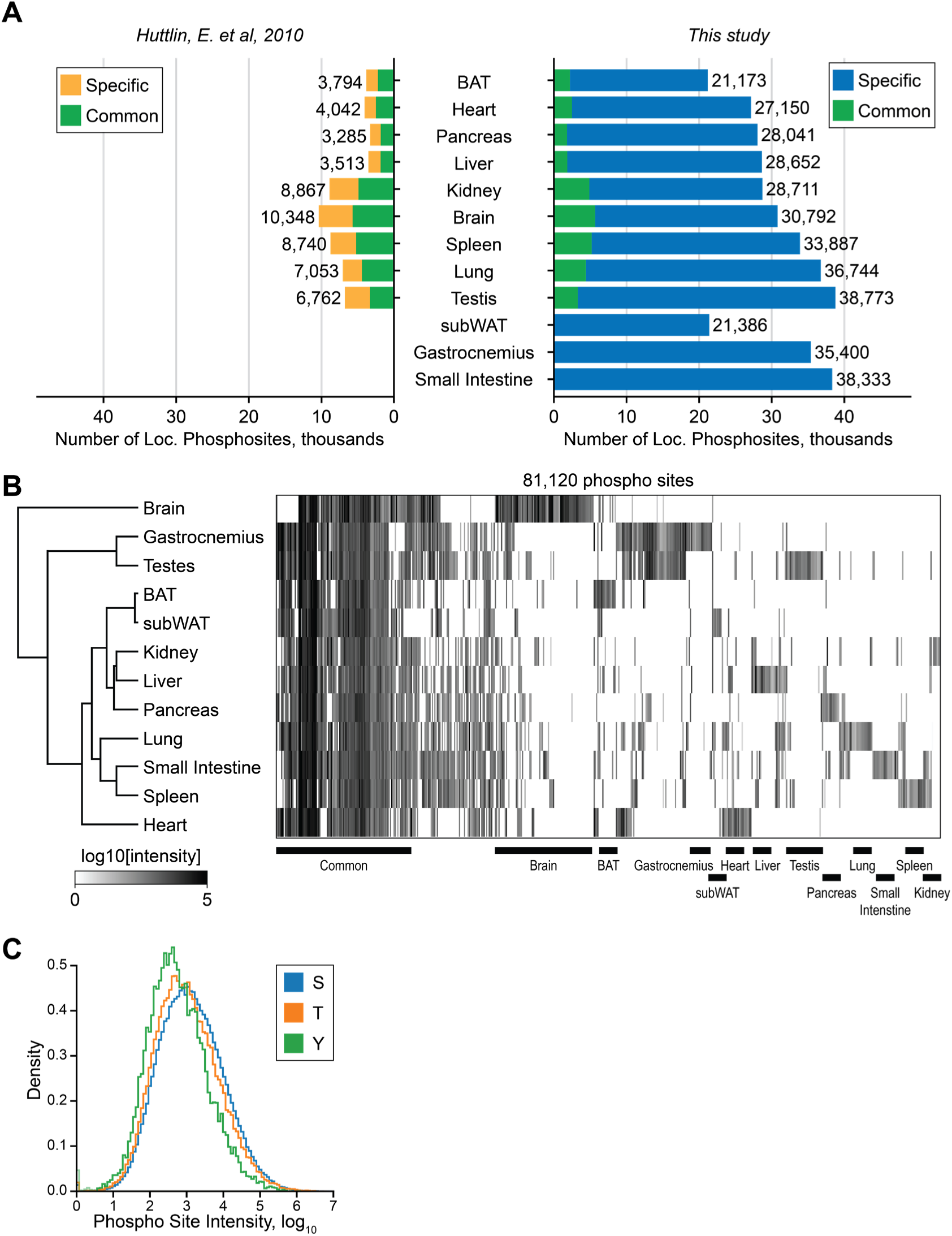
Mouse Phosphorylation Atlas Comparison to Huttlin et al. (A) Overlap of phosphorylation sites detected in this study and by Huttlin et al. (B) Heatmap showing the relative intensities of phosphosites detected across tissues. Note that there are many phosphorylation sites with similar intensities across all tissues, but each tissue tends to have clusters of phosphosites that are uniquely high intensity (C) Normalized distribution of phosphorylation site intensities for S, T, and Y localizations.

**Supplementary Figure 6.**
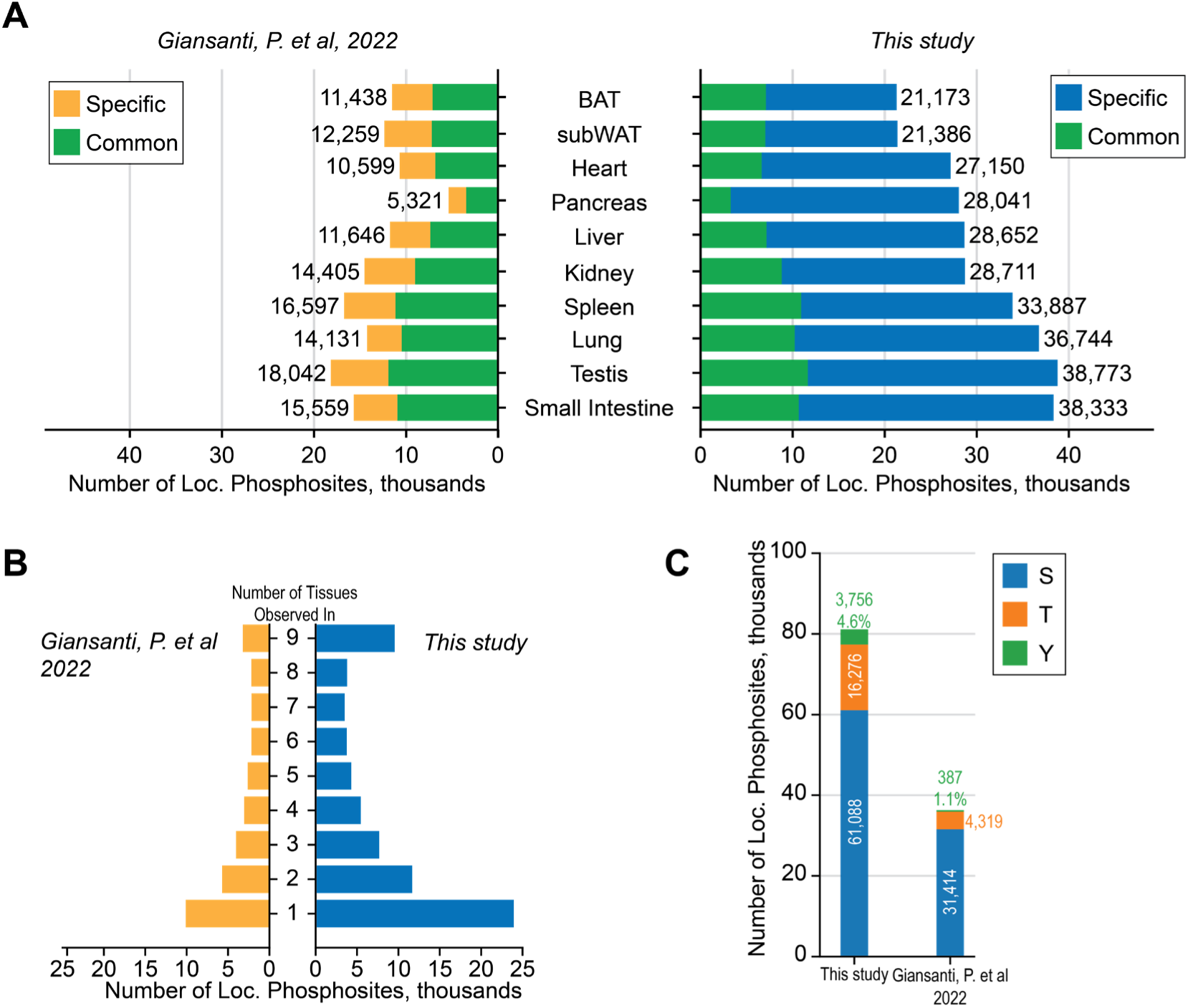
Mouse Phosphorylation Atlas Comparison to Giansanti et al. (A) Overlap of phosphorylation sites detected in this study and by Giansanti et al. (B) Tissue Specificity of Detected Phosphorylation sites, excluding the pancreas, brain, and gastrocnemius tissues from comparison. The y-axis indicates the number of tissues in which a phosphosite was detected. (C) The total number of S, T, Y phosphorylation sites are shown for this study and Giansanti et al. The percentage of the total sites occurring on phosphotyrosine residues is indicated for both studies.

**Supplementary Figure 7.**
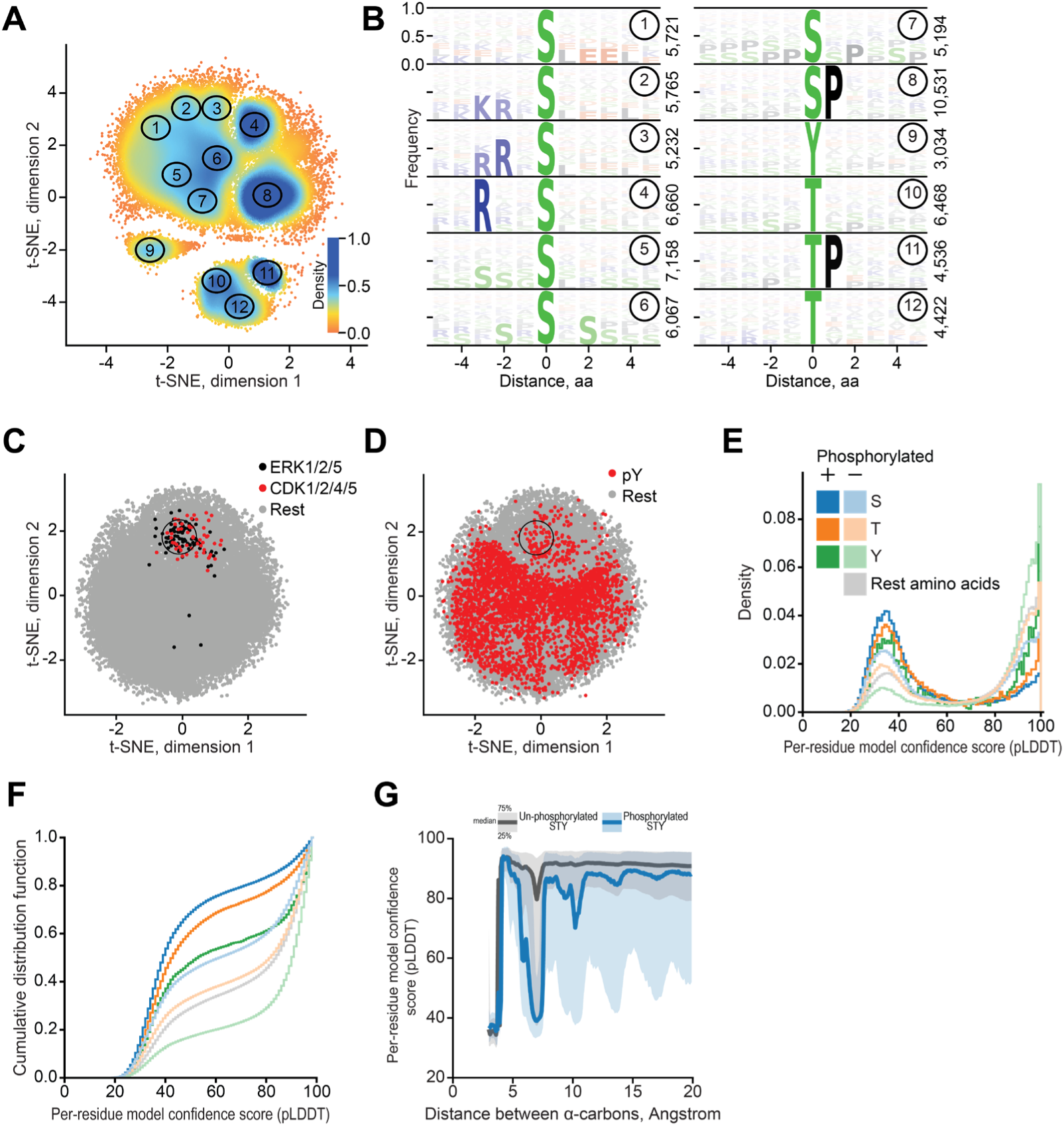
Sequence and Structural Context of Phosphosites. (A) Two-dimensional representation of all phosphosites and their 5-amino acid flanking sequences, including the central amino acid in the comparison. Each cluster has been manually selected to highlight the densest regions. (B) Sequence logo plot for all clusters depicted in (A). (C) Similar to Figure 5A with highlights of ERK and CDK kinases based on the PhosphoSitePlus database. (D) Similar to Figure 5A with highlights of all phosphorylated tyrosine sites. (E) Distribution of structural confidence scores for phosphorylated (+) and unphosphorylated STY sites. (F) Similar to (E) but presented as a cumulative distribution plot. (G) Confidence score distribution based on the distance between alpha-carbons surrounding phosphorylation site and phosphorylation state – phosphorylated and un-phosphorylated S/T/Y.

**Supplementary Figure 8.**
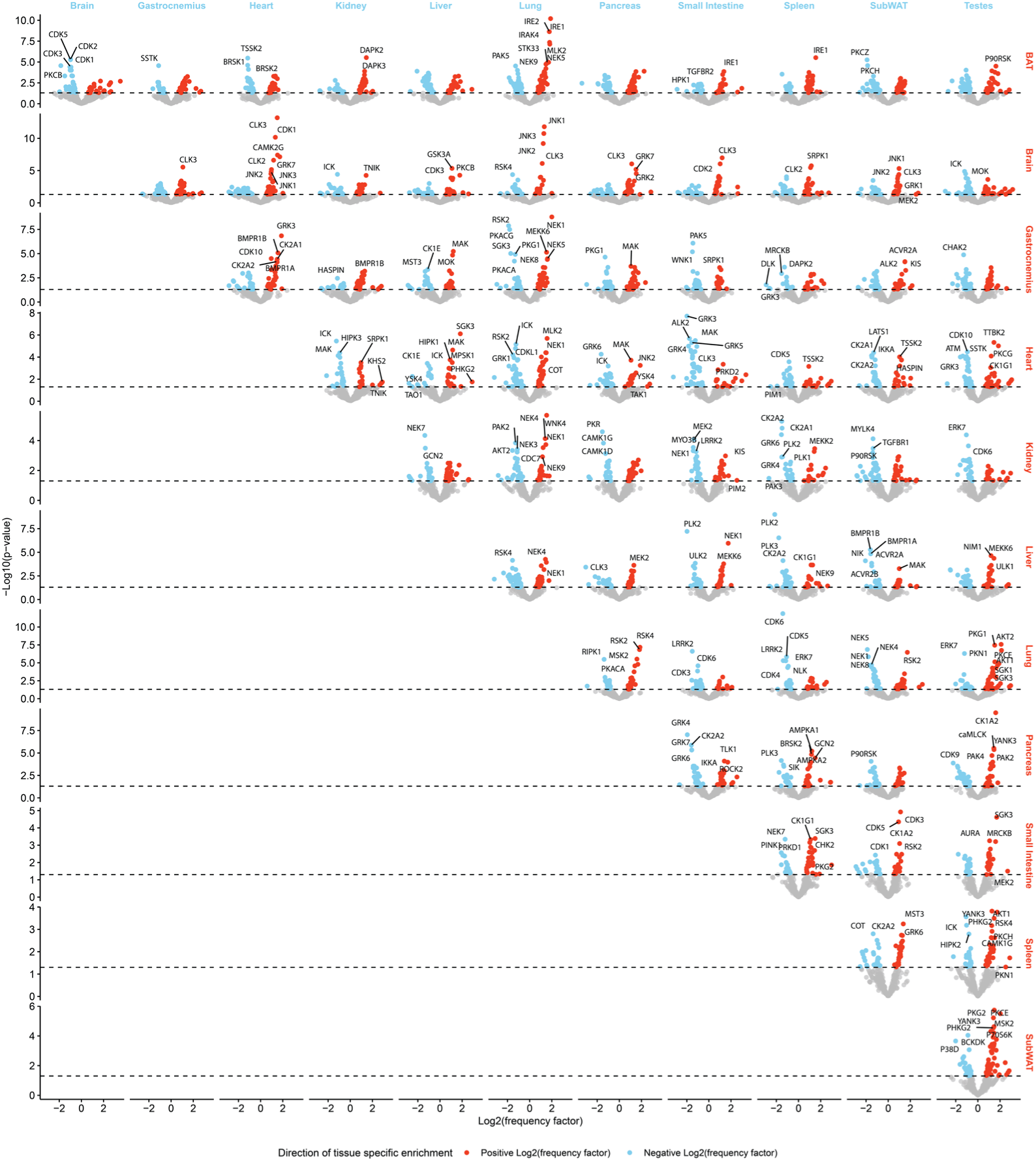
Kinase Enrichment Across Tissues. For each possible tissue-tissue pair, the top tissue-specific phosphorylation sites were selected to perform a kinase motif enrichment analysis. The frequency of how often a kinase was predicted to act on a site in a tissue was plotted on the x-axis. The p-value is shown on the y-axis. The red dots indicate significant kinases in the tissues that are labelled in red on the right side of the plot. The blue dots correspond to the tissues plotted across the top of the graph. Source data is provided in **Supplementary Table 5**.

**Supplementary Figure 9.**
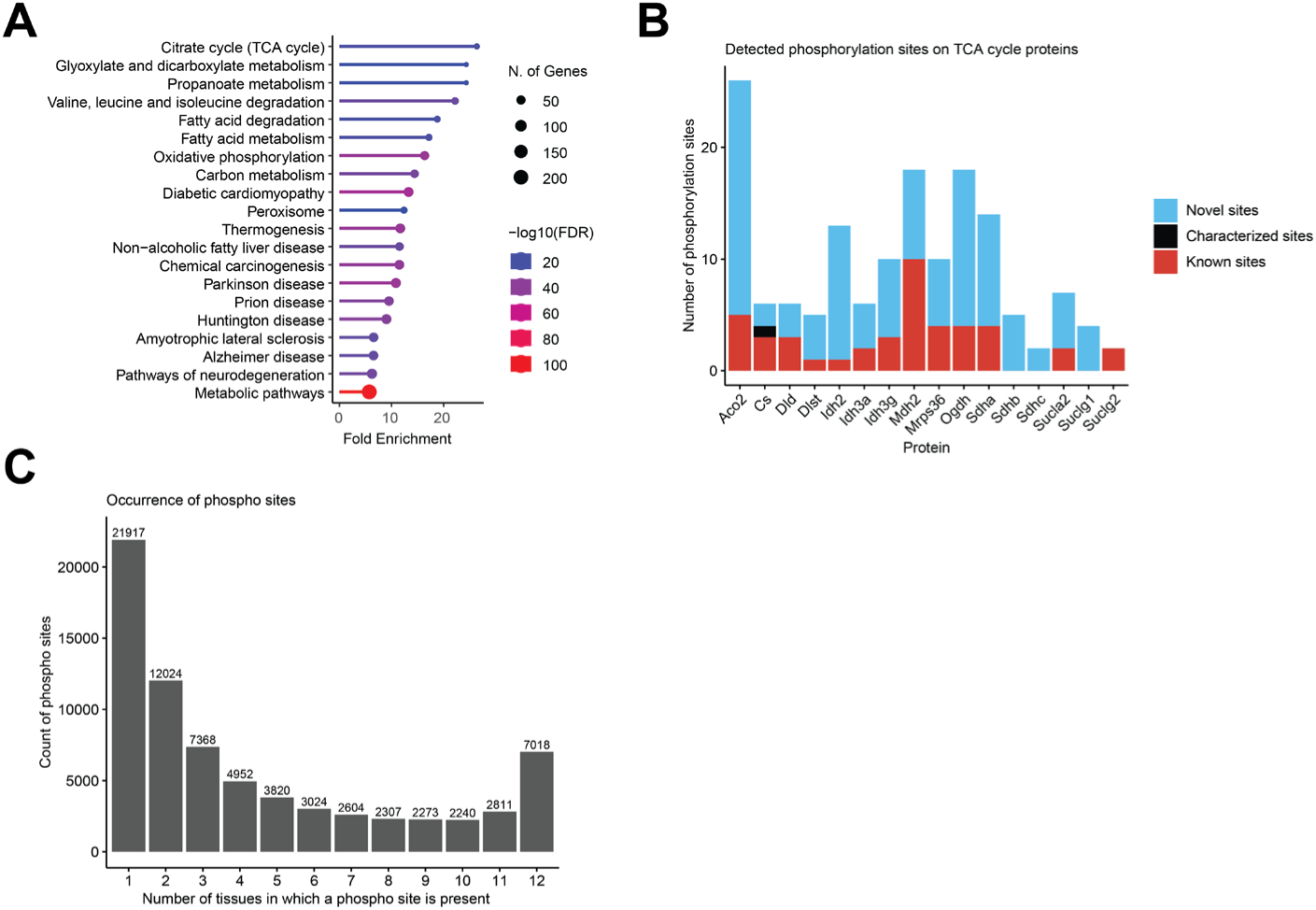
TCA cycle phosphorylation events and occurrence of phosphorylation sites across tissues. (A) GSEA using the KEGG pathway database, FDR 0.05, minimal pathway size 2, through the ShinyGO app.^83^ Input: Proteins whose phosphorylation sites showed an absolute z-score above 2. (B) Number of phosphorylation sites detected on tricarboxylic cycle proteins (subset based on MitoCarta 3.0 pathways). Sites that occur in multiple tissues are summed. Characterized sites according to PhosphoSitePlus are also known sites and are only counted in the characterized sites category. (C) Bar plot indicating the number of phosphorylation sites that occur in a specific number of tissues.

**Supplementary Figure 10.**
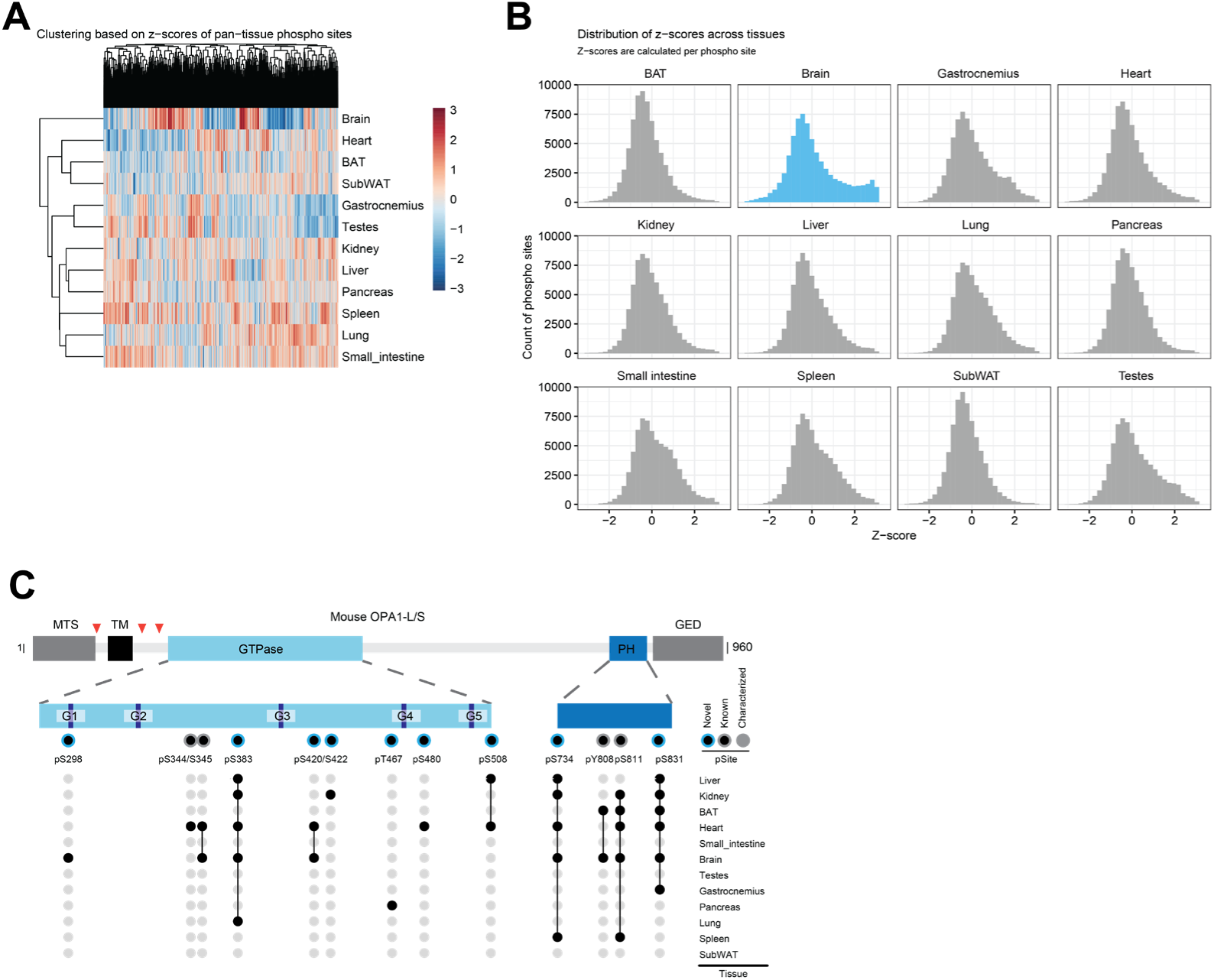
Z-score based outlier analysis and phosphorylation sites on Opa1. (A) Heatmap of z-scores calculated across all tissues. Only sites that occur in all tissues were considered. (B) Distribution of phosphorylation site z-scores in each tissue, using the entire imputed dataset. (C) Protein domain representation of mouse OPA1 as shown in Figure 5D with additional phosphorylation sites not observed in brain tissue. The occurrence of OPA1 phosphorylation site within all twelve dataset is shown below.

**Supplementary Table 1.** Phosphopeptide Standards Summary and Dilution Series Data

**Supplementary Table 2.** Summary of all detected phosphosites for the EGF-stimulation experiments.

**Supplementary Table 3.**Summary of all detected phosphosites from the phosphoproteomics experiment for mouse tissues.

**Supplementary Table 4.** Summary of all protein groups detected in the proteomics experiment for mouse tissues.

**Supplementary Table 5.** Summary of kinase enrichment analysis for mouse tissues.

**Supplementary Table 6.** Expression enrichment and mitochondrial annotation of phosphosites.

