## Supplemental Text for "Fast and Deep Phosphoproteome Analysis with the Orbitrap Astral Mass Spectrometer"

MS1 Orbitrap Scan Settings

Because Astral MS2 acquisition can be performed in parallel with the Orbitrap MS1 scan, we selected the 240,000 resolving power at 200 *m/z* scan for high-resolution MS1 scan acquisition. This scan takes just over 0.5s[1], so we selected a 0.6s cycle time (the instrument control software allows cycles times in 0.1s increments) to maximize the Orbitrap MS1 duty cycle.

MS2 Maximum Inject Time

The timing sequence for Astral analysis has been previously described by Stewart *et al*.[2] Using a maximum injection time of 3.5 ms ensures that the ion accumulation time is close to maximally parallelized with other steps in the scan sequence. This setting enables a scan rate of ~200 Hz to be maintained throughout the runs. Below, the $MS^1$ and $MS^2$ cycle times are plotted for our 15-minute active gradient method with a 2 Th DIA window iterating over a 380-980 m/z range (a total of 300 windows). Cycle times of ~0.6 and ~1.5s are observed for the $MS^1$ and $MS^2$ scans, respectively (**Supplemental Figure 1E**). Maintaining short cycle times is essential as the columns employed in this study yield median chromatographic peak widths less than 8 seconds for our 15-minute method (as shown in **Supplemental Figure 1B**).

DIA *m/z* Range

The *m/z* range iterated over during DIA analysis is a key method parameter. The size of this range limits the achievable cycle time (or the minimum isolation window width). In our experiments, we utilized a 380-980 *m/z* range, a common range used for DIA proteomics.[2–4] Enriched phosphopeptide samples could exhibit different *m/z* distributions than proteomics samples due to the mass shift from phosphorylation and increased missed cleavages[5] so we compared the use of a 380-980 *m/z* range with a 480-1080 *m/z* range for the analysis of EGF-stimulated HeLa phosphopeptides. We observe that the use of the 480-1080 *m/z* range decreases the number of phosphosites by ~6% (**Supplemental Figure 2D**), validating our use of a 380-980 *m/z* range in this study. However, we note that the optimal *m/z* range likely depends on the sample type being analyzed.

Astral AGC target

The AGC target for the Astral analyzer influences the number of charges that are targeted for each scan. We utilized a target of 5e4 (500%) for DIA phosphoproteomics. We compared this to a method using a target of 1e4 for analysis of the EGF-stimulated HeLa phosphopeptide samples. We note that there is no considerable difference in the depth achieved with these two methods (**Supplemental Figure 2E**). We note that the MS2 scan total ion current distributions are identical for both methods (**Supplemental Figure 2F**). This observation is explained by looking at the MS2 inject time distribution, with that both methods reach the maximum injection time setting of 3.5 ms for most scans (**Supplemental Figure 2G**). The absence of an effect on phosphoproteomic depth then arises from the AGC target not being reached in either method for these particular analyses.
